# Developmental tuning of functional manifold dimensionality across the human brain

**DOI:** 10.64898/2026.07.24.740635

**Authors:** Erica L. Busch, Nicholas B. Turk-Browne

**Affiliations:** Department of Psychology Wu Tsai Institute, Yale University

## Abstract

Neural representations vary in their complexity across brain regions and tasks. How this variation emerges over human development remains poorly understood. We estimated intrinsic dimensionality in five naturalistic fMRI datasets (*N* = 781 unique participants, aged 3 months to 53 years) with T-PHATE — a nonlinear manifold learning method robust to noisy, autocorrelated signals. In adults, brain regions relevant to a task had higher-dimensional activity than task-irrelevant regions across auditory, visual, and audiovisual stimuli. The modulation of representational complexity by tasks was absent in infants, emerged in early childhood, and strengthened logarithmically through adolescence. It reflected a selective collapse in dimensionality in task-irrelevant regions, relative to a resting-state baseline, rather than an expansion of dimensionality in task-relevant regions. Such compression followed a trajectory from global and nonselective in infants to local and precise by adulthood. These results identify selective compression as a developmental engine of functional specialization: rather than adding complexity where it is needed, the brain dynamically pares it away where it is not.

## Introduction

The human brain must form rich representations to support complex cognition while operating efficiently within biological constraints. One way of exploring this tension is by tracking the intrinsic dimensionality of brain activity — the minimum number of variables needed to adequately capture the underlying structure and complexity of population-level activity patterns [1–3]. Intrinsic dimensionality offers a lens into the computational complexity of neural processing, where higher dimensionality reflects more independent processing modes and greater representational capacity and flexibility [4–9]. Accordingly, higher-dimensional activity has been linked to efficient learning and cognitive processing in adults, suggesting dimensionality is not merely a descriptive property but a functional resource the brain actively deploys [10–13].

Much of what we know about the computational complexity of neural activity comes from electrophysiological recordings of individual brain regions during constrained laboratory tasks in animals [14–17], or from neuroimaging studies focused on specific brain networks and static, controlled paradigms in adult humans [10, 12, 18–20]. This leaves two key questions unanswered. First, how is intrinsic dimensionality distributed and modulated across the human brain during rich, naturalistic cognitive processes — engaging perception, memory, affect, social inference, etc. — typical of human mental life? Second, how do these large-scale dimensional properties of human brain function emerge over development, from infancy through adulthood?

We address these questions by quantifying intrinsic dimensionality across the whole brain during naturalistic tasks (story listening and movie watching) and during baseline rest and sleep states in 781 participants aged 3 months to 53 years from five open-access fMRI datasets. We introduce an intrinsic dimensionality (ID) estimator grounded in the T-PHATE manifold learning framework [21]. T-PHATE was designed to handle the high noise, nonlinear geometry, and temporal autocorrelation inherent to fMRI data — features that cause standard dimensionality reduction methods to overestimate signal complexity [22–24].

We find that task-relevant brain regions maintain higher dimensionality than task-irrelevant regions in adults, that this functional specialization is absent in infancy then develops logarithmically from early childhood through adolescence, and that task engagement drives selective compression in task-irrelevant regions relative to rest/sleep. Together, these results reveal that the brain allocates computational complexity selectively — maintaining representational richness where it is needed while compressing activity elsewhere — and that this capacity develops gradually over the first two decades of life.

## Results

### T-PHATE accurately estimates intrinsic dimensionality from noisy, simulated signals

We estimated intrinsic dimensionality (ID) from fMRI timeseries using T-PHATE [21] as the number of eigenmodes required to explain 90% of diffusion variance (Figure 1A). We validated this estimator on five synthetic manifolds with known ID and benchmarked it against five existing ID methods spanning linear covariance (principal components analysis [PCA], local PCA with participation ratio) to nonlinear estimators designed for complex, high-dimensional data (maximum likelihood estimation [MLE], minimum neighbor distance – maximum likelihood [MiND_ML], dimensionality from angle and norm concentration [DANCo]; see Table S1) [25]. To test robustness to realistic timeseries noise, we added to each manifold noise generated using a first-order autoregressive process with varying autocorrelation strength and magnitude [22]. Accuracy was assessed using a normalized error rate 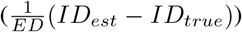, which accounts for the different ground-truth dimensionalities (*ID*_*true*_) and embedding dimensions (*ED*) across datasets.

**Figure 1:**
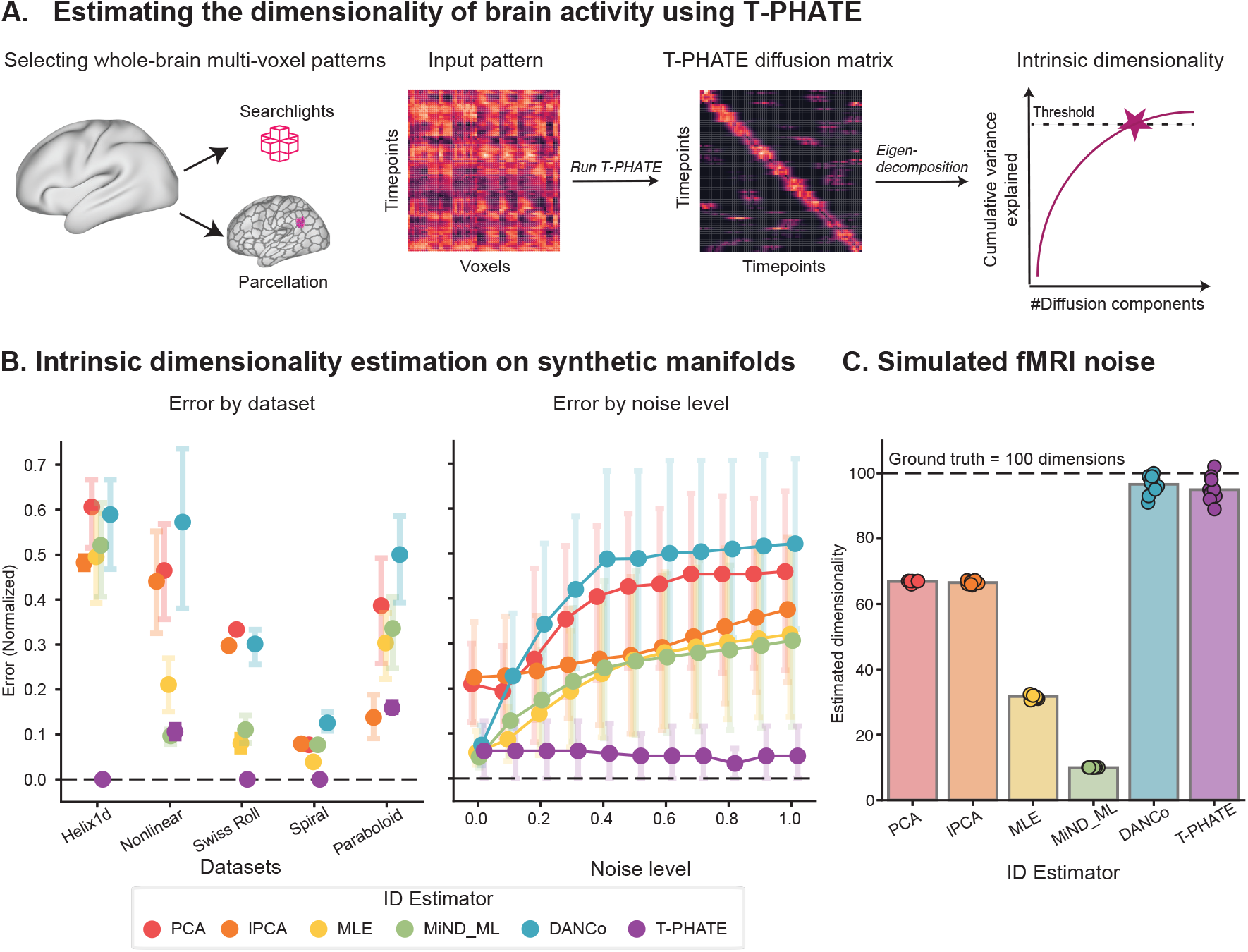
Protocol and validation of T-PHATE intrinsic dimensionality estimator. (A) Intrinsic dimensionality (ID) estimation began by selecting multi-voxel activity patterns. Searchlights allow for an ID estimate at each brain voxel based on its neighborhood of surrounding voxels (within a 5-voxel radius). Parcellations divide the brain into non-overlapping regions (e.g., Schaefer 400 parcellation; [26]). T-PHATE uses the multi-voxel timeseries data to compute a samples × samples diffusion operator representing the probability of taking a random walk between data points along a manifold, weighted both by manifold geometry estimated with the PHATE algorithm [27] and the temporal autocorrelation between samples [21]. Eigendecomposition of the T-PHATE diffusion operator yields the variance explained by each diffusion eigenvector. Setting a threshold for the target amount of variance explained (90%) results in an estimate of the data manifold’s ID. (B) Evaluating the performance of T-PHATE ID estimation against benchmark ID estimators on synthetic manifold datasets with varying magnitudes of noise. (left) T-PHATE estimations achieved lower error, exactly estimating the true manifold dimensionality in three of the five datasets and being among the closest for the other two datasets. (right) T-PHATE performance was robust to noise, while other estimators become significantly less accurate at higher noise levels. (both) Points show mean normalized error, averaged across all autocorrelation strengths (*α* ∈ [0, 1)) and across all noise levels (left panel) or all datasets (right panel); error bars show the 95% CI of the mean. (C) ID was estimated on 10 simulated timeseries datasets with no low-dimensional structure (i.e., true ID= 100 dimensions). Points represent simulated datasets, bars show means.

T-PHATE showed the lowest error overall and greatest robustness to noise across all synthetic manifolds, exactly recovering true ID for three of the five datasets across all noise levels (Figure 1B). Nonlinear methods (MLE, MiND_ML, DANCo) matched T-PHATE performance at lower noise levels but degraded substantially with increased noise, while linear methods (PCA, lPCA) showed consistently higher error across datasets. To assess whether estimators would accurately report high ID, we tested all estimators on 10 simulated fMRI datasets with no latent structure. Simulated datasets had 200 samples and 100 statistically uncorrelated features (mean inter-feature *r* = − 0.001 *±* 0.001; one-tailed *t*(9) = − 2.56, *p* = 0.99). T-PHATE and DANCo most accurately recovered high ground-truth dimensionality of 100 dimensions (Figure 1C). Together, these results demonstrate that T-PHATE is unique in its both robust and accurate estimation of ID from nonlinear, noisy, autocorrelated signals characteristic of fMRI.

### Higher dimensional activity associated with reliable responses to auditory narratives

Having validated our ID estimator, we first characterized dimensionality in adults during naturalistic information processing. We analyzed the Narratives dataset [28], in which 40 adults each listened to four audio-only, first-person narrative stories during fMRI (Table S2). To quantify how engaged different brain areas were with the stimulus, we computed intersubject correlation (ISC) within parcels of the Schaefer 400 atlas [26]. ISC is a model-free way of gauging task-relevant brain regions during rich and temporally dependent stimuli (like narratives or movies) by evaluating the consistency of a brain area’s responses across participants, so that parcels with higher ISC are more reliably engaged in processing stimulus content [29, 30]. We estimated the ID of activity in each parcel using T-PHATE, which allowed us to compare the task-relevance and the dimensionality of activity patterns across the brain, within each participant and story.

Participants showed the most reliable stimulus-driven activation (i.e., highest ISC) in auditory cortex, temporoparietal junction (TPJ), and prefrontal cortex (PFC; Figure 2A). ID maps demonstrated a similar spatial pattern: areas with higher ISC (particularly TPJ and PFC) also showed higher ID relative to the rest of cortex (Figure 2B). ISC and ID maps showed highly consistent spatial patterns across stories (ISC: mean pairwise *ρ* = 0.99; ID: mean pairwise *ρ* = 0.92; Figure 2C). Further, spatial patterns of whole-brain ISC and ID were significantly correlated within participant, even across stimuli (mean *ρ* = 0.353, all *p* < 0.0001; Figure 2C), suggesting that more reliable task-evoked brain responses were also higher dimensional. To confirm the specificity of this effect, we performed a spatial permutation test (10,000 iterations), shuffling ISC scores across parcels relative to ID scores to generate a null distribution of *ρ* values (within participant and story); we computed z-scores of the true *ρ* relative to this null distribution. Observed ISC-ID z-scores were significantly greater than expected by chance for all four stories (mean *±* SD; *z* = 4.546 *±* 2.576, all *p* < 0.0001; Figure 2D), confirming a robust relationship between the task engagement and dimensionality of a brain region during naturalistic listening.

**Figure 2:**
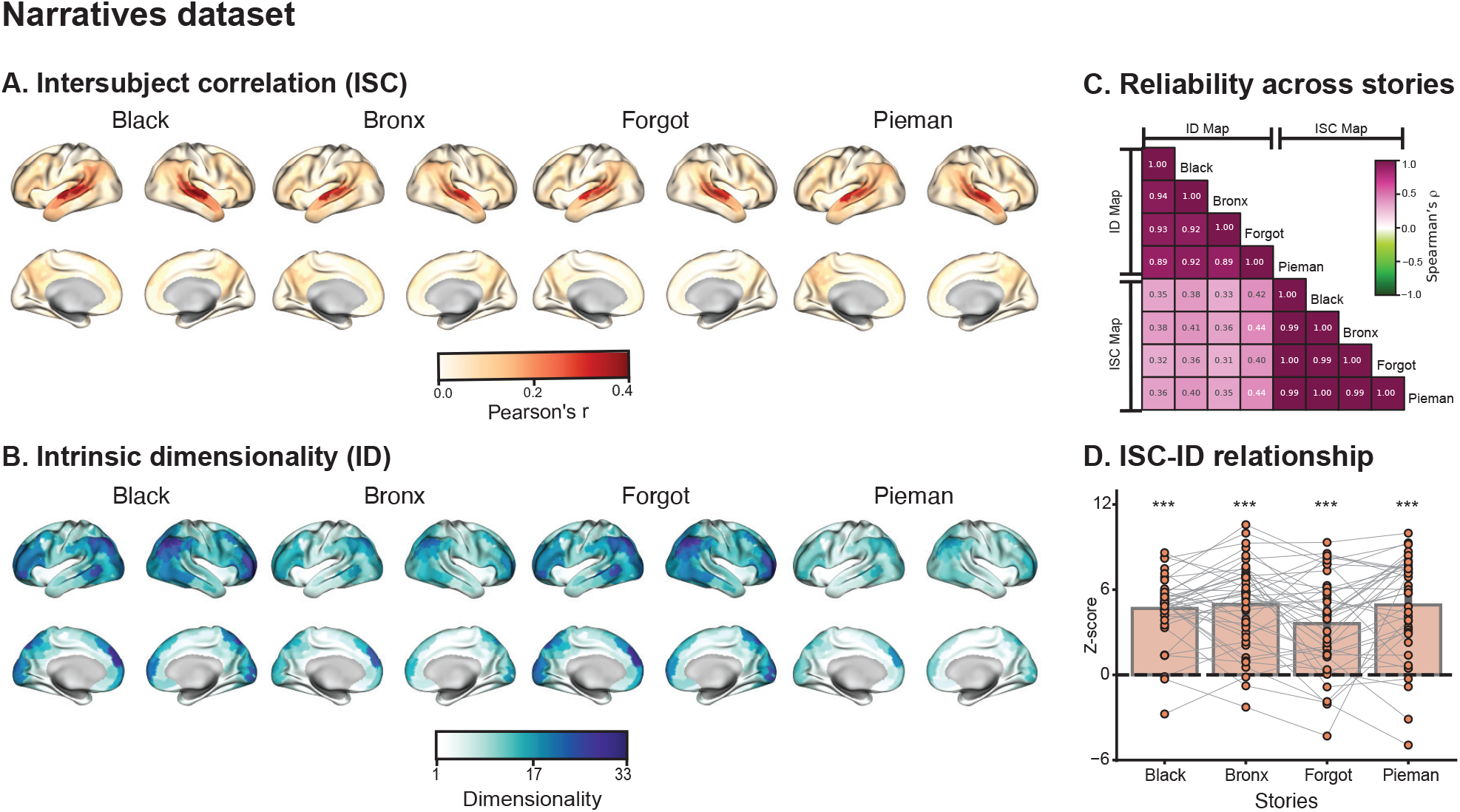
Narratives dataset: Adults listening to audio-only stories. (A) ISC maps averaged across participants for the four stories. (B) ID maps averaged across participants for the four stories. (C) Spearman’s *ρ* between all pairs of parcel-wise ID and ISC brain maps, averaged across participants. The near-diagnonal blocks (upper left, lower right) show within-metric, across-story (ID-ID, ISC-ISC) correlations within-participant, averaged across participants. The off-diagonal block (lower left) shows cross-metric (ISC-ID) correlations within-participant, averaged across participants. ISC-ID map correspondence within story for each participant, quantified as *ρ* converted to z-scores using spatial permutation tests (10,000 iterations). Bars represent the average *z* across participants, points represent individual participants, and lines connect the same participant across stories. *** *p* < 0.001 (one-sample t-test, two-tailed).

### Task-relevant dimensionality in adults generalizes to the visual modality

To test whether the observed ISC-ID relationship extends beyond the auditory domain, we analyzed adults (*N* = 12, mean age = 21.4 *±* 3.4 years) from the Rest/Movie dataset who watched a silent cartoon movie (“Aeronaut”) during fMRI. We computed ISC and ID using whole-brain overlapping searchlights (5-voxel radius sphere) rather than parcels, since there are no infant-specific parcellations that would allow for direct comparison with adults. As in the Narratives dataset, brain regions showing the highest ISC — now concentrated in the visual cortex — also exhibited the highest dimensionality (Figure 3A,B, adults). Within-participant ISC-ID correlation z-scores were significantly positive (mean *z* = 6.66, *t*(11) = 4.56, *p* = 0.001), demonstrating that the principle of higher dimensionality in task-relevant regions generalizes from auditory to visual stimuli (Figure 3C). These results provide converging support for the conclusion that task-relevant regions in the adult brain maintain higher-dimensional representations than task-irrelevant regions.

**Figure 3:**
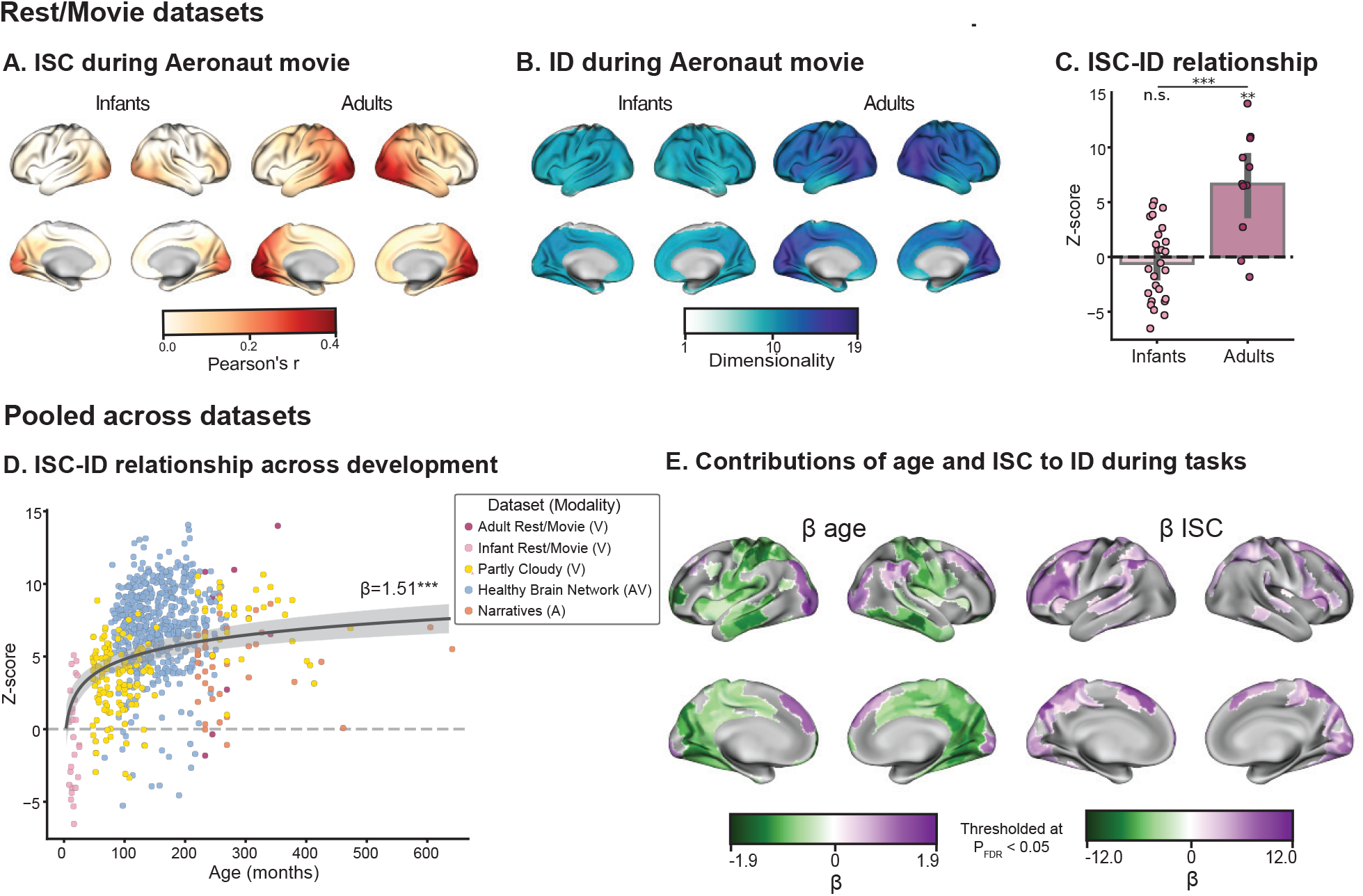
ISC-ID correspondence during naturalistic tasks across development. (A) ISC searchlight maps averaged across infant (*N* = 26) and adult (*N* = 12) participants who watched the movie “Aeronaut”. (B) ID maps computed in the same manner. (C) Correlation of the ISC and ID searchlight maps for each participant, averaged across participants. True Spearman’s *ρ* values were converted to z-scores using a permutation test (10,000 iterations). (D) Relationship between the z-scored ISC-ID correlation and log-transformed age for each participant, pooled across datasets (colors). Line shows a mixed-effects model fit predicting participants’ z-scores from age (controlling for head motion [FD], sex, and dataset) which revealed a significant logarithmic increase in ISC-ID correlation with age (*β*_*age*_ = 1.51, *p* < 0.001). Distinct contributions to a parcel’s ID by participant age (left) or the parcel’s ISC (right), measured using mixed-effects models at each parcel (controlling for FD, sex, and dataset). *β* coefficients are thresholded at *p*_*FDR*_ < 0.05. Purple indicates regions where the predictor is associated with higher ID; green indicates association with lower ID. ** *p* < 0.01, *** *p* < 0.001

### Dimensional specificity emerges during childhood

Having discovered a relationship between stimulus processing and dimensionality in adults, we next examined how this correspondence emerges developmentally. We first analyzed infants (*N* = 26, mean age = 10.9 *±* 4.8 months) from the Rest/Movie dataset who watched the same movie as adults (Aeronaut). The overall magnitude of ISC was lower among infants than adults (mean *±* SD; infants: *r* = 0.033 *±* 0.072; adults: *r* = 0.133 *±* 0.097; Welch’s *t*(34.49) = − 3.72, *p* = 0.0007), but followed a similar pattern of higher ISC in visual cortex, confirming reliable stimulus processing (Figure 3A). However, infant ID was uniformly low and less variable across the entire cortex (mean *±* SD; 9.31 *±* 1.67), contrasting with adults, who showed higher and more variable ID, with peaks in visual regions 11.46 *±* 2.98; *t*(19.24) = − 2.27, *p* = 0.035; Figure 3B). Critically, the robust positive correlation between ISC-ID in adults (from earlier: mean *z* = 6.66, *t*(11) = 4.56, *p* = 0.001) was absent in infants (mean *z* = − 0.60, *t*(25) = − 0.89, *p* = 0.38); the difference in correlation between groups was highly significant (*t*(14.46) = 4.51, *p* = 0.0004; Figure 3C).

To track how the task specificity of ID develops between infancy and adulthood, we analyzed two additional datasets spanning broader age ranges. The Partly Cloudy dataset included children aged 3-12 years (*N* = 122) and adults (*N* = 33) watching a silent cartoon movie. The Healthy Brain Network (HBN) dataset included children and adolescents aged 5-21 years (*N* = 528) watching an audiovisual cartoon movie (see Figure S1 and Table S2 for age breakdown). Both datasets showed the expected distribution of ISC across the brain, with highest ISC in visual, posterior parietal, and somatosensory cortices for Partly Cloudy and visual and auditory cortices for HBN (Figures S2A, S3A). Pooling across all five datasets (*N* = 765, age range 3 months – 53 years, we confirmed that mean whole-brain ISC increases as a function of age (mixed-effects model accounting for head motion [mean framewise displacement, FD], sex, and dataset): *β*_*age*_ = 0.015, 95% *CI* = [0.008, 0.021], *z* = 4.17, *p* = 0.00003; Figure S4A), consistent with the infant-adult difference observed above.

Despite the increase in ISC across development, mean whole-brain ID decreased slightly across the pooled age range (mixed-effects model accounting for mean FD, sex, and dataset): *β*_*age*_ = − 0.32, 95% *CI* = [− 0.626, 0.004], *z* = − 1.99, *p* = 0.047; Figure S4B). This finding indicates that the developmental trajectory of task-driven ID is more nuanced than an overall expansion with age, as was shown in the infant-adult comparisons, and points to more interactive, regional trends which may be washed out in whole-brain averaging [31, 32].

To model developmental change in the correspondence of ISC and ID continuously across the full age range, we pooled each participant’s z-scored correlation of ISC and ID brain maps from all five datasets and predicted ISC-ID z-scores from age (again accounting for FD, sex, and dataset). Given prior work showing nonlinear growth trajectories in neurodevelopmental data, we tested linear, logarithmic, and quadratic age terms. A log-transformed age term provided the best fit (*AIC* = 3709.97, *BIC* = 3742.38) compared to linear (increase in AIC/BIC relative to log-transformed AIC/BIC: Δ*AIC* = 6.37, Δ*BIC* = 6.37) and quadratic (Δ*AIC* = 5.38, Δ*BIC* = 10.01) alternatives, consistent with prior work [33]. The logarithmic model revealed a significant positive effect of age on ISC-ID z-scores (*β*_*age*_ = 1.51, 95% *CI* = [1.03, 2.00], *z* = 6.10, *p* < 0.001; Figure 3D), confirming that the task specificity of dimensionality increases steeply in early childhood and plateaus through adolescence into adulthood.

### Age and tasks make distinct contributions to regional dimensionality

The developmental increase in ISC-ID coupling could reflect spatially dynamic, age-related changes in ISC, ID, or both. To tease apart these contributions and identify brain regions driving dimensionality changes, we modeled how log-transformed age and ISC predicted ID in each brain parcel, pooling across all five datasets (*N* = 765 unique participants). For each parcel, we used mixed-effects models to predict ID from log-transformed age and ISC, controlling for sex, FD, and dataset; we retained coefficients surviving FDR correction (*p*_*FDR*_ < 0.05; Figure 3E).

ID increased with age in visual cortex, right TPJ, and parts of medial PFC (38 of 400 parcels), while decreasing in somatomotor, parietal, and medial temporal regions (203 parcels). In contrast, ID tracked ISC positively in visual cortex, parts of auditory cortex, lateral PFC, and precuneus (155 parcels); no parcels showed a significant negative relationship between ISC and ID. Moreover, no parcels showed a significant interaction effect, indicating that maturation and task engagement shape dimensionality through parallel mechanisms: maturation enables more efficient computation (dampening complexity where possible), while task engagement is associated with representational richness (maintaining or expanding where necessary) [7, 11, 34, 35]. On its own, however, task-related dimensionality dynamics could reflect elevated dimensionality in task-relevant regions, suppressed dimensionality in task-irrelevant regions, or both. Adjudicating between these requires a task-free reference, which we introduce below by comparing task states against rest.

### Dimensionality of task-irrelevant regions collapses over development

To understand how task states shape dimensionality, we compared ID during rest (or natural sleep in infants) with ID during movie-viewing in the Rest/Movie and HBN datasets. These datasets were uniquely useful for this analysis because they contained both types of data.

In the Rest/Movie dataset, the adults (*N* = 12) each completed resting and movie-viewing runs, allowing for within-participant comparisons, whereas separate groups of infants completed sleeping (*N* = 20, mean age *±* SD, = 11.2 *±* 5.0 months) and movie-viewing runs (*N* = 26, 10.9 *±* 4.8 months). Resting and sleeping ID exceeded movie-viewing ID in both age groups (paired samples t-test in adult cohort: *t*(11) = 3.11, *p* = 0.011; independent samples t-test in infant cohort: *t*(31.76) = 3.98, *p* = 0.0004; Figure 4A,B).

**Figure 4:**
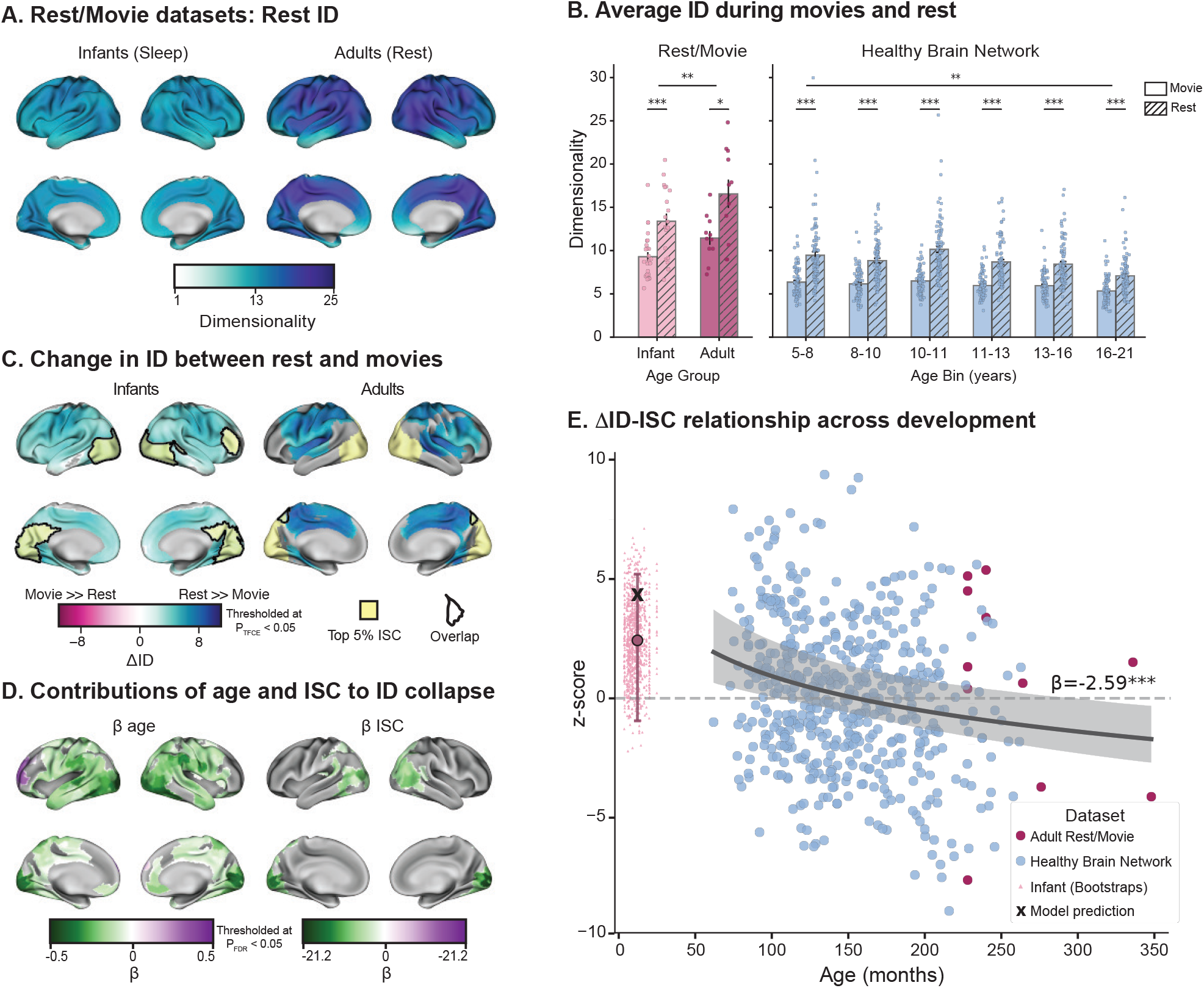
Specificity of collapse in ID for movie-watching versus rest or sleep. (A) ID during natural sleep (infants) and rest with fixation (adults) averaged across participants. (B) ID for movie-viewing and rest/sleep in Rest/Movie adults and infants (left) and HBN children/adolescents (right). HBN participants were binned into approximately age groups of equal size for visualization purposes. All groups show significantly higher ID during rest/sleep than movie-viewing. (C) Rest/Movie dataset ΔID was computed as resting/sleeping ID minus movie-watching ID (related samples for adults, independent samples for infants). Significance was assessed using FSL’s randomise with threshold-free cluster enhancement (TFCE) correction, and only searchlights surviving correction at *p*_*TFCE*_ < 0.05 were visualized. Searchlights showing the top 5% of ISC values within group are overlaid in yellow on the ΔID maps for each group. Clusters where the top 5% of ISC searchlights (yellow) overlap with searchlights showing significant ΔID are outlined in black. (D) Relative contributions of age and ISC to ΔID were assessed at each parcel using mixed effects modeling, controlling for sex, FD, and dataset. Data for this analysis was pooled across the Rest/Movie adults and infants and HBN datasets. *β* coefficients for log-transformed age (left) and ISC values (right) were visualized for each parcel in the brain, thresholded at *p*_*FDR*_ < 0.05. (E) The relationship between ID collapse and stimulus-related activity was quantified as the Spearman correlation between a participant’s ΔID map (i.e., rest ID minus movie ID) and ISC map, converted to a z-score via permutation testing. The relationship between ΔID-ISC z-scores and development was assessed using mixed effects modeling to account for sex, FD, and dataset. Line shows logarithmic model fit (*β*_*age*_ = − 2.59, *p* < 0.0001) for the adult Rest/Movie and HBN datasets, which allowed for within-subject comparison across resting state and movie-watching scans; band shows the 95% CI of model fit. Since infants in the Rest/Movie dataset did not have within-subject repeated sleep/movie-watching scans, bootstrap resampling was used to simulate a distribution of infant ΔID-ISC z-scores. True participant ΔID-ISC z-scores are shown as circular points, the bootstrapping distribution of infant scores shown as triangles, the mean and 95% CI of the bootstrap distribution shown as a point and error band, and the model’s predicted predicted z-score for the mean-age infants is shown as **‘X’**.

In the HBN dataset (*N* = 528), a linear model predicting ID from task, age, and their interaction (controlling for FD and sex) confirmed a significant effect of task (rest > movie: *β*_*task*_ = 3.91, *t*(1049) = 8.01, *p* < 0.001), a marginal negative effect of age (*β*_*age*_ = −0.004, *t*(1049) = −1.92, *p* = 0.065), and a significant negative *age* × *task* interaction (*β*_*age* × *task*_ = − 0.008, *t*(1049) = − 2.68, *p* = 0.007). As with the average movie-viewing ID results above, this model indicates that the overall magnitude of collapse in dimensionality induced by tasks decreases over development and suggests nuanced specificity with regard to which regions show these effects (Figure S3C,D).

To understand the spatial distribution of dimensionality attenuation across task states, we computed ΔID maps by subtracting movie-watching ID maps from resting state ID maps. In adults, ID decreased significantly from rest to movie in parietal, frontal, and motor cortices — notably, searchlights with the lowest ISC during movie viewing — while visual cortex, where ISC was strongest, showed no significant ΔID. In infants, ID decreased globally across the brain from sleep to movie without regional specificity. The overlap between task-relevant searchlights (i.e., high ISC) and searchlights with significant ΔID was quantified with a Szymkiewicz-Simpson overlap coefficient. This confirmed that among adults, task-driven searchlights didn’t demonstrate dimensionality collapse (overlap coefficient = 0.053), but among infants, dimensionality collapse occurred brain-wide — even among visual regions with highest ISC (overlap coefficient = 0.937). In other words, task engagement causes selective dimensional collapse in regions less relevant for processing a visual movie (as indexed by lower ISC) in adults but a nonselective collapse in infants. Widening the age range with the HBN dataset revealed that the regional specificity of dimensionality collapse as observed in adults (but not infants) emerged in late childhood (Figure S3D).

To disentangle the contributions of development and task relevance on ΔID, we pooled data across the infant and adult Rest/Movie datasets with the HBN dataset and used mixed-effects modeling to predict how each parcel’s ID changed across states depending upon the participant’s age and the parcel’s degree of task relevance (controlling for FD, sex, and dataset). Age predicted decreased ΔID (or greater dimensionality preservation) in visual, auditory, temporal, and posterior parietal cortices, while ISC predicted decreased ΔID in an overlapping set of regions including visual and posterior parietal cortices and TPJ (Figure 4D). In other words, age predicted a smaller drop in ID in most of the brain, and the regions with stronger task-relevant activity also showed smaller ID drops. No regions showed a significant *age* × *ISC* interaction, indicating that age and ISC operate in parallel to predict preservation of dimensionality during task engagement.

Finally, to quantify the brain-wide relationship between task-relevant activity and dimensionality collapse between rest and task across development, we computed the Spearman correlation between participants’ ΔID maps and their ISC maps in the adult Rest/Movie and HBN datasets. These scores were then converted to z-scores via spatial permutation testing (10,000 iterations). Finally, a mixed-effects model was used to predict ISC-ΔID z-scores from age (accounting for mean FD, sex, and dataset) (*β*_*age*_ = − 2.59, 95% *CI* = [− 3.485, − 1.697], *z* = − 5.68, *p* < 0.0001, Figure 4E). This analysis demonstrated that, with age, dimensional collapse becomes more negatively correlated with stimulus-driven responses. In other words, younger children show non-selective dimensionality collapse during tasks relative to rest, but with age, the collapse becomes more specific to regions which are less task-relevant, and more task-relevant areas demonstrate reliable preservation of dimensionality. The adult Rest/Movie and HBN datasets afforded within-participant computation of the ISC-ΔID z-scores, as all participants completed both resting state and movie-watching scans. However, infant participants did not have both kinds of data. To test the fit of this model on the youngest participants, we computed a bootstrapping distribution using resampling to approximate a mean and confidence interval of the estimated ISC-ΔID effect among infants. The model was then applied on the average infant age, and the model’s prediction was compared with the bootstrapping distribution. The model-predicted z-score for mean age fell within the bootstrapped confidence interval, showing that the model’s validity extended to the infants.

Together, these findings show that offline states (rest/sleep) maintain high dimensionality across cortex, and that online states (movies/stories) constrain activity to lower-dimensional subspaces. This constraint is initially global and non-selective in infancy and early childhood and becomes increasingly specific to task-irrelevant brain regions with development. By adulthood, the brain selectively preserves dimensional complexity in task-relevant regions while collapsing it elsewhere.

## Discussion

Neural populations process and encode the features of simple, well-defined stimuli by expanding or compressing their modes of information processing [9, 36, 37]. We examined such shifts in dimensionality over the whole brain, in response to naturalistic tasks, and across human development. We leveraged existing large-scale fMRI datasets spanning infancy to adulthood that presented participants with spoken narratives or movies to engage low- and high-level perception, social cognition, memory, affect, and other cognitive systems. The use of whole-brain measurements allowed us to characterize fluctuations in dimensionality across the entire cortex.

Our findings suggest several organizing principles: First, task-relevant brain regions have higher-dimensional representations than task-irrelevant regions. Second, this difference is absent in infancy, emerges in early childhood, and strengthens through adolescence. Third, ID is high and uniform across brain regions during rest. Fourth, the difference in dimensionality between task-relevant and task-irrelevant regions reflects a decrease from rest for task-irrelevant rather than an increase for task-relevant. Fifth, this collapse of dimensionality in task-irrelevant regions also undergoes developmental refinement.

The robust coupling between ISC and ID further confirms that regions that reliably process stimulus content maintain richer and more complex representational geometries [38–41]. This held across sensory modalities: auditory cortex during audio-only narratives, visual cortex during visual-only movies, and both during audiovisual movies. These results suggest an additional way to think about functional specialization beyond selective activation, in terms of the computational complexity with which a region represents information [7, 8, 42]. The consistently high dimensionality across tasks in multimodal convergence zones like the TPJ and posterior parietal cortex may reflect the role of these regions in integrating information across the sensory, cognitive, and social domains common to naturalistic stimuli [28, 43–45].

Among the most striking findings in our study is the developmental trajectory of the ISC-ID relationship. Infants had reliable ISC in visual cortex but uniformly low and spatially non-specific ID. By early childhood, a positive ISC-ID relationship emerged and strengthened logarithmically through development, consistent with progressive functional specialization of cortical computations [31, 33, 46]. Region-based modeling revealed independent contributions of age and task engagement (operationalized as ISC) to this trajectory. We interpret the age-related decrease in ID across much of cortex as computational refinement or attunement, where older brains perform less-specialized computations more efficiently [35, 47–49]. On the other hand, sensory cortices and multimodal hubs like TPJ maintain or expand dimensionality with age, preserving representational capacity where required for specialized processing demands [11, 32, 42]. ISC-related increases in ID, orthogonal to age, may reflect enhanced task processing through the deployment higher-dimensional representations. The absence of an age-by-ISC interaction suggests that these may be separable mechanisms.

Using fMRI data collected during awake rest or natural sleep, we found that the resting brain establishes a ceiling on ID, reflecting the unconstrained nature of activity at rest. Task engagement consistently suppressed dimensionality below resting levels across all age groups, extending findings from animal models to the human brain during naturalistic tasks [37, 50]. During tasks, adults showed selective suppression in task-irrelevant (i.e., low ISC) regions while preserving high dimensionality in task-driven (i.e., high ISC) regions — reflecting efficient, targeted allocation of computational resources [17, 34]. In contrast, infants showed global, non-specific ID collapse, suggesting that early attentional engagement broadly suppresses neural complexity without functional precision. Importantly, global dimensionality collapse in the infant brain included regions that were reliably stimulus-responsive (i.e., visual cortex during movie-viewing) — providing reassurance that this effect does not solely reflect noise or inattention. The developmental shift from global to selective parallels the maturation of executive function and top-down attentional control, following a logarithmic developmental trajectory consistent with other neurodevelopmental trajectories [33].

Our results are qualified by several limitations. Because naturalistic stimuli engage a rich and interrelated set of cognitive processes simultaneously, it is difficult to attribute individual dimensions to specific stimulus features or cognitive functions. This is a tradeoff inherent to using more ecologically valid and dynamic stimuli, with the upside of allowing the collection of similar data across developmental participants from infants to adults. Accordingly, we focused on changes in relative dimensionality across brain regions, tasks, and ages, rather than absolute values. Further, more research will be needed into the computational changes that accompany shifts in ID. Other geometric properties of the neural manifold — including curvature, topology, and the organization of representational subspaces — may undergo fine-tuning even after ID appears stable [5, 6]. Finally, understanding the neurobiological mechanisms underlying dimensional changes — including changes in neural tuning, excitation/inhibition balance, population size, and connectivity — as well as the trait- and state-level psychological correlates of dimensional shifts, remain open questions that will require linking fMRI with structural and behavioral measures [13, 18]. The current work paves the way for future multi-modal analyses spanning feature modeling, behavior tracking, and learning interventions to understand neurocomputational maturation across the lifespan.

## Materials and Methods

### Data

We used five fMRI datasets spanning infancy through adulthood to examine intrinsic dimensionality across development and naturalistic sensory contexts, including auditory, visual, audiovisual, and offline (resting or sleeping) states. Basic information about the datasets, tasks, participants, and ages is summarized in Table S2, and age distributions are shown in Figure S1. Two classes of simulated datasets were additionally used for benchmarking.

### Narratives dataset

The Narratives dataset includes adult participants listening to auditory stories while undergoing fMRI scanning [28]. We analyzed a subset of 40 participants (*N* = 29 female, mean age = 23.5 *±* 7.9 years) who each listened to four stimuli (personal narratives from “The Moth Radio Hour”: “Bronx”, “Pie Man (PNI)”, “Black”, and “Forgot”) presented in separate runs. Participants reported normal hearing, no neurological disorders, and provided written informed consent. Data were collected at Princeton University with institutional approval and informed consent. Data were obtained from OpenNeuro (ds002345). Full acquisition and preprocessing details are provided in the Supplemental Methods.

### Rest/Movie dataset

#### Adults

Adult participants (*N* = 12, mean age = 21.4 *±* 3.4 years, *n* = 7 female) completed both resting fixation scans and viewed a silent cartoon movie (“Aeronaut”). Data were collected at Yale University with institutional approval and informed consent. The movie stimulus “Aeronaut” is a silent 3-minute long segment of an audiovisual short film called “Soar” (https://vimeo.com/148198462). The video extended 45.5 degrees of visual angle wide and 22.5 high. The order of presentation between rest and movie-watching runs was counterbalanced across participants.

#### Infants

Infant data included sleeping scans (*N* = 20 sessions from 14 unique infants; mean age = 11.2 *±* 5.0 months) and movie-watching scans (*N* = 26 sessions from 18 unique infants; mean age = 10.9 *±* 4.8 months). Sleep state was determined by behavioral monitoring during scanning. Data were collected at Yale University and Princeton University with institutional approval and parental consent. These data are openly available via DataDryad (https://doi.org/10.5061/dryad.nvx0k6dzf) and were used in a previous publication [51]. Detailed acquisition and preprocessing procedures are described in the Supplemental Methods.

### Partly Cloudy dataset

The Partly Cloudy dataset includes *N* = 122 children (3.5°12 years) and *N* = 33 adults (18°39 years) who watched a silent 5.6-minute cartoon film “Partly Cloudy” during fMRI scanning [45] (pixar.com/partly-cloudy#partly-cloudy-1). This film emphasizes social cognition, pain perception, and theory of mind among non-human characters. Data were collected at MIT with institutional approval and informed consent. They are publicly available via OpenNeuro (ds000228). Acquisition and preprocessing details are provided in the Supplemental Methods.

### Healthy Brain Network dataset

We analyzed a subset of participants from the Healthy Brain Network Biobank [52] who completed both resting state and movie-watching scans. The movie-watching task involved watching “The Present”, an emotionally evocative short audiovisual animated movie (approximately 4 minutes) focusing on the relationship between a boy and a dog. The final sample included 528 participants (mean age = 12.32 *±* 3.5 years). Data were collected across three separate imaging centers and accessed via data.healthybrainnetwork.org. Detailed exclusion criteria, acquisition parameters, and preprocessing pipelines are described in the Supplemental Methods.

### Simulated datasets

We evaluated dimensionality estimation using synthetic nonlinear manifolds with known ground-truth dimensionality from the scikit-dimension package [25, 53]. Additional null datasets were generated from high-dimensional noise with no latent structure, to assess estimator behavior under realistic timeseries noise conditions.

### Benchmark manifolds

We tested the T-PHATE, PCA, lPCA, MiND_ML, and DANCo methods on five benchmark nonlinear manifolds with known ground-truth dimensionality. Key information on each dataset is included in Table S1. The “M5a_Helix1d” dataset represents a one-dimensional helical curve embedded in three-dimensional space, forming a spiral trajectory that tests the algorithm’s ability to recover low-dimensional structure from curved embeddings. The “M6_Nonlinear” manifold consists of a highly nonlinearly six-dimensional manifold embedded in a thirty-six-dimensional space. The “M7_Roll” dataset, commonly known as the “Swiss roll,” is a two-dimensional rectangular sheet that has been rolled into a spiral shape in three-dimensional space, representing a classic test case for manifold learning algorithms. The “M13b_Spiral” manifold forms a one-dimensional helical surface with varying curvature and density embedded in a thirteen-dimensional space, challenging dimensionality estimators to distinguish signal from the complex embedding geometry. Finally, the “Mp1_Paraboloid” represents a three-dimensional paraboloid nonlinearly embedded in twelve-dimensional space according to a multivariate Burr distribution (*α* = 1). These benchmark datasets span intrinsic dimensionalities between one and six, with varying degrees of curvature, nonlinearity, and geometric complexity, providing a rigorous test bed for evaluating the accuracy and robustness of our diffusion-based dimensionality estimation approach.

The noise generation process begins by adding independent Gaussian noise to the first timepoint. For each subsequent timepoint, the noise is generated as a weighted combination of the previous timepoint’s noise and new independent Gaussian noise, controlled by an autocorrelation coefficient *α*. This creates a first-order autoregressive (AR(1)) noise process where the strength of temporal correlation is determined by *α*, with higher values (*α* → 1) producing stronger autocorrelation and lower values (*α* → 0) producing nearly independent noise across timepoints.

Formally, given input data *X* ∈ ℝ ^(*T* ×*N*)^, where *T* is the number of timepoints and *N* is the number of features, the noisy data 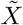 is generated as:

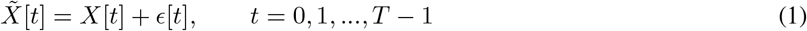

The noise terms follow a stationary autoregressive process:

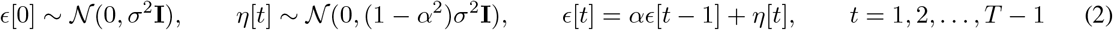

Where *ϵ*[0] initializes the process by drawing from the stationary distribution, *η*[*t*] represents independent Gaussian noise at each timepoint with variance scaled by (1 − *α*^2^) to ensure stationarity, *σ* ∈ [0, 1] controls the noise magnitude, *α* ∈ [0, 1) determines the autocorrelation strength, and **I** denotes the *N* × *N* identity matrix. Together, *σ* and *α* vary independently, such that the marginal variance of *ϵ*[*t*] equals *σ*^2^ at all timepoints regardless of *α*.

### Simulated null data

We used 10 simulated timeseries datasets (200 samples, 100 features) with no low-dimensional structure (i.e., features were optimized to be independent from one another). Data were generated using a package optimized to control the correlational structure of datasets with realistic noise and temporal properties [54]. Thus, we could assess whether the estimators could recover the true 100-dimensionality from these datasets or were prone to mis-estimating dimensionality in the face of noise.

### Whole-brain approaches

Intrinsic dimensionality was computed across the brain using both volumetric searchlights and cortical parcellations. Searchlights (5-voxel radius, 1331 voxels/searchlight) were used for Infant and Adult Rest/Movie datasets to avoid reliance on adult-defined areal boundaries. Voxels were considered as searchlight centers if all participants within a group/task (e.g., all infants viewing Aeronaut) contained that voxel in their whole-brain mask, and if > 80% of voxels in the searchlight had non-zero values in a given scan. All voxels that met these criteria were used as the center of the searchlight. For older cohorts (with greater sample sizes per dataset), dimensionality was also estimated within parcels from the Schaefer-400 atlas [26]. These complementary approaches yielded spatially resolved dimensionality maps for each participant and task. For alignment to standard spaces and visualization of these parcellations on the cortical surface, we used the `neuromaps` and `brainspace` packages [55, 56].

### Intersubject correlation

Stimulus-driven reliability was assessed using spatial intersubject correlation (ISC). For each brain area, multi-voxel timeseries were vectorized and correlated between each participant and the average of all other participants. For each searchlight or parcel, we extracted a multi-voxel activity pattern of timepoints *t* by voxels *v*, which was then vectorized, resulting in one *t* ∗ *v* vector **x** per participant. Then, the ISC score for a participant *i* at a given searchlight or voxel was the Pearson correlation between *x*_*i*_ and the average of all the other participants’ *x* vectors. This procedure is repeated, holding out each participant’s data and computing 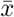 across the remaining participants, and across all spatial locations in the brain (i.e., each searchlight or parcel). This produced participant-specific ISC maps reflecting shared stimulus-evoked processing [29, 30].

### Intrinsic dimensionality analysis

For each participant, scan, and brain area, preprocessed fMRI data were represented as a matrix **X** ∈ ℝ ^*T* ×*V*^ , where rows correspond to timepoints and columns correspond to voxels. Intrinsic dimensionality was estimated using the T-PHATE framework described below.

#### Intrinsic dimensionality estimation from T-PHATE

To estimate intrinsic dimensionality while accounting for both nonlinear geometry and temporal structure in fMRI data, we applied T-PHATE (Temporal Potential of Heat-diffusion for Affinity-based Transition Embedding), a manifold learning algorithm specifically designed for neural timeseries data [21]. T-PHATE extends PHATE [27] by integrating two views of the data: a geometry-driven affinity structure (“PHATE view”) and an explicit temporal autocorrelation model (“Temporal view”), which are combined into a unified diffusion operator.

The steps to getting the T-PHATE diffusion operator from input data **X** work as follows (with steps 1 through 6 coming from the PHATE algorithm):

1. Compute pairwise distances between data points, to measure how dissimilar every pair of data points is and provide an initial geometry of the dataset.

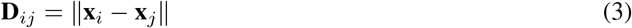
2. Convert distances to affinities via an adaptive kernel. The locally adaptive bandwidth *σ*_*i*_ is derived from the distance to the *k*-nearest neighbor of **x**_*i*_ and affords handling regions of varying density.

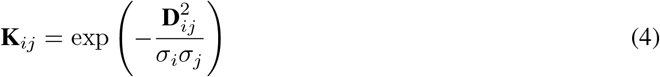
3. Row-normalize the affinity matrix to obtain a Markov transition matrix, describing the probability of moving from one data point to another in a random walk.

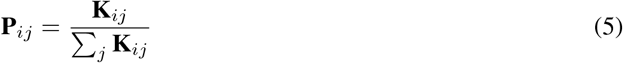
4. Raise **P** to power *t*, simulating a random walk over the data graph. The diffusion time parameter *t* controls the scale of structure captured, allowing PHATE to integrate local and global relationships.

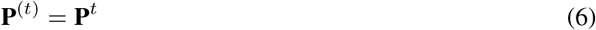
5. Compute potential representations by taking the negative logarithm. This stabilizes diffusion differences, emphasizing meaningful differences between data points while reducing noise.

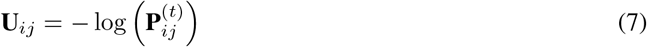
6. Compute distances between the potential profiles of each data point, capturing the intrinsic manifold geometry.

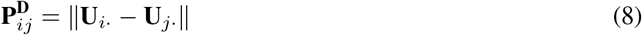
7. From the original input data **X**, take the temporal mean of each voxel:

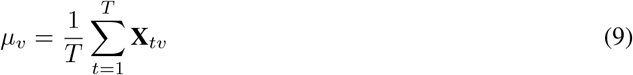
8. For each time lag *τ* ∈ {0, 1, …, *T* − 1}, obtain the covariance between the timeseries and a lagged copy of itself to define the autocovariance of **X**_·,*v*_ as:

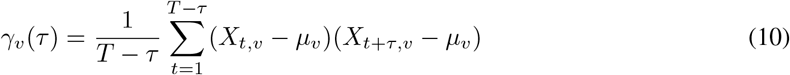
9. Average the normalized autocovariance vectors across voxels to get a single autocorrelation vector **c**:

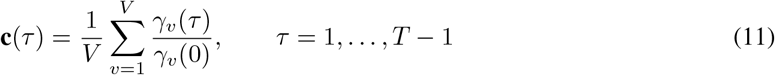
10. Smooth **c** with a rolling average over *w* samples to serve as a damping tool for possible jittering around where

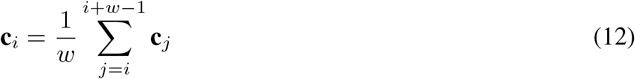
11. Find the first lag *τ* where **c** = 0, which defines the maximum width of smoothing for the temporal affinity matrix **A**.

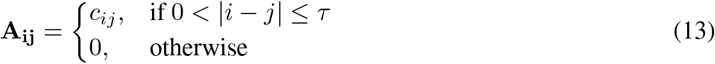
12. Row-normalize **A**:

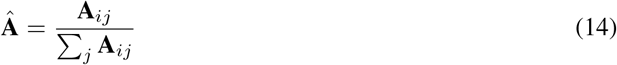
13. Raise **Â** to the power of *t* as in Equation 6 and multiply with **P**^**D**^ from Equation 8 to combine the temporal view with the PHATE view to get the T-PHATE diffusion operator **P**:

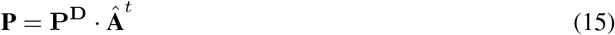
14. Compute the eigendecomposition of **P**, the row-stochastic diffusion operator constructed from T-PHATE, as:

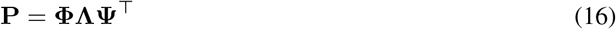

**P** is generally nonsymmetric, so the eigendecomposition is comprised of right and left eigenvectors (**Φ, Ψ**) and eigenvalues (Λ) ordered such that *λ*_1_ ≥ *λ*_2_ ≥ . . . ≥ *λ*_*T*_ . Thus, the eigenvectors represent the principal modes of diffusion on the manifold, while the eigenvalues quantify the contribution of each mode to the overall diffusion process. Accordingly, the largest eigenvalues *λ*_*i*_ → 1 correspond to persistent and slowly-decaying diffusion modes and the large-scale geometric structure in the manifold, whereas the smaller eigenvalues *λ*_*i*_ → 0 correspond to local fluctuations and noise. The number of significant eigenvalues reflects the intrinsic dimensionality of the diffusion process.
15. The cumulative proportion of explained variance for each eigenvalue then becomes:

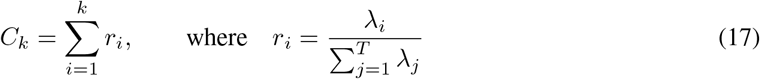

The intrinsic dimensionality (ID) of the input data is estimated as the minimum number of eigenmodes required to capture a specified threshold proportion *θ* of the total variance: *ID* = min *k* : *C*_*k*_ ≥ *θ*. In the present study, we used *θ* = 0.9, or a 90% variance explained threshold.

### Benchmark methods

To evaluate T-PHATE, we compared it against established intrinsic dimensionality estimators implemented in `scikit-dimension` [25], including PCA, local PCA, maximum likelihood estimation (MLE), MiND_*ML*_, and DANCo. These methods estimate dimensionality using covariance structure, nearest-neighbor distance distributions, or geometric concentration properties. Full mathematical formulations are provided in the original publications [57–59] and included in Supplemental Methods.

### Statistical analyses

Correspondence between ISC and ID within task and participant were assessed by computing the true Spearman correlation *ρ* between parcel-wise ISC and ID maps. To assess the spatial specificity of this effect, we generated a null distribution by permuting a participant’s ISC values relative to their ID values before recomputing the *ρ* values, and then computed the z-score of the true *ρ* relative to the null distribution. We assessed statistical significance of z-scores relative to zero using one-sample t-tests with two-tailed p-values. Comparisons across tasks within age groups were performed using paired samples t-tests. Comparisons across groups were performed using independent samples Welch’s t-tests (for unequal variances). Age group and task effects were assessed using ordinary least squares regression, including covariates for sex and head motion (mean framewise displacement [FD] per scan). Mixed effects models were used when performing analyses across pooled datasets, including fixed effects for sex and FD and random intercepts for datasets. 95% confidence intervals were computed using bootstrap resampling (10,000 iterations). Regression models were implemented using Python’s *statsmodels* package (version 0.14.4).

Statistics for parcel-wise brain maps were computed using bootstrap resampling (10,000 iterations) and thresholded using FDR correction (*p*_*FDR*_ < 0.05). Statistics for searchlight brain maps were computed using FSL’s randomise function (paired samples for adults, independent samples for infants), corrected for multiple comparisons using threshold-free cluster enhancement (TFCE), and visualized where *p*_*TFCE*_ < 0.05.

### Code availability

Data analysis code are written as custom Python scripts (version 3.9.19) and available at the following github repository: https://github.com/ericabusch/task_dim. T-PHATE (version 1.2.1) is available as a Python package at https://github.com/KrishnaswamyLab/TPHATE and can be installed via pip https://pypi.org/project/TPHATE [60].

### Generative AI statement

During the preparation of this work, the author(s) used Claude Code for assistance with programming data visualization code. The author(s) reviewed and edited the output as needed and take full responsibility for the content of the published article.

## Acknowledgments

E.L.B. was supported by an NSF Graduate Research Fellowship (award no. 2139841). N.B.T.-B. was supported by the NIH (grant no. R01MH069456), NSF (grant no. 1839308) and CIFAR. Additional internal funding was provided by the Faculty of Arts and Sciences and the Wu Tsai Institute at Yale University. We thank Ping Luo of the Wu Tsai Institute for assistance with high-performance computing, Tristan Yates and Smita Krishnaswamy for helpful conversations, and Marvin Chun, Samuel McDougle, and Arielle Baskin-Sommers for their feedback on earlier versions of this work.

## Supplemental Materials

### Supplemental Methods

#### MRI Acquisition and Preprocessing

Detailed acquisition parameters and preprocessing pipelines are summarized below for each dataset.

#### Narratives dataset

##### MRI data acquisition

Functional and anatomical images were collected on a 3T Siemens Magnetom Prisma scanner using a 64-channel head coil. Whole-brain functional images were acquired in an interleaved fashion (48 slices per volume, 2.5mm isotropic resolution) using a gradient-echo EPI (repetition time [TR] = 1.5 s; echo time [TE] = 31 ms; flip angle = 67) with a multiband acceleration factor of 3 and no in-plane acceleration. Each stimulus was collected in a separate run, resulting in the following volumes for “Black” (550), “Bronx” (390), “Forgot” (574), “PiemanPNI” (294).

##### MRI data processing

We used preprocessed data from [28], accessed via Datalad (OpenNeuro database accession number ds002345). Briefly, fMRIprep was used to perform coregistration, slice-time correction, and nonlinear alignment to the MNI152 template brain [61]. Timeseries data were detrended with regressors for motion (translation and rotation in x, y, z dimensions), white matter, and cerebrospinal fluid before smoothing with a 6 mm full-width-half-max (FWHM) Gaussian kernel. Full information about data acquisition and preprocessing can be found in [28].

#### Rest/Movie dataset

##### Adult participants

The adult fMRI data includes 12 participants (*N* = 7 female, mean age = 21.4 *±* 3.4 years) who completed both an awake resting state scan (staring at fixation) and viewed a silent cartoon movie. Resting-fixation scans were 5 minutes, during which participants were instructed to stare at a white fixation cross (presented at 2 visual degrees) without thinking about anything in particular. The movie stimulus “Aeronaut” is a 3-minute long segment of an audiovisual short film called “Soar” (https://vimeo.com/148198462). The video was presented at 45.5 visual degrees in width and 22.5 visual degrees in height and was played without audio in the scanner. The order of presentation between rest and movie-watching runs was counterbalanced across adult participants (Table S2). Adults were recruited from the New Haven, Connecticut community and data were collected at the Brain Imaging Center in the Faculty of Arts and Sciences at Yale University. The study was approved by the Human Subjects Committee (HSC) at Yale University, and all adults provided informed written consent.

##### Infant participants

The infant fMRI data includes 20 sleeping scans (*N* = 14 unique infants, *N* = 7 female, mean age = 11.2 *±* 5.0 months, range = 3.9 − 24.9 months) and 26 movie-watching scans (*N* = 18 unique infants, *N* = 14 female, mean age = 10.9 *±* 4.8 months, range = 3.6 − 20.1 months) (Table S2; Fig. SS1). Of the 14 unique infants with sleeping fMRI data, four infants completed 2 sleeping scans each, and one infant completed 3 sleeping scans. One infant completed more than one sleeping scan during the same session; the remainder were spread across multiple sessions (mean = 5.3 months between consecutive sessions, range = 1.3 − 15.0 months). Sleeping scans were collected from infants who fell asleep naturally in the scanner. Sleep state was assumed based on extended eye closure and stillness while in the scanner, as observed online via an MR-safe camera, and researchers turned off the visual display. Infants slept in the scanner for variable durations (mean = 4.3 *±* 1.3 min, range = 1.8 − 6.0 minutes). Participants in the Aeronaut task watched the same silent 3-minute long segment as did the adult participants. Of the 18 unique infants with movie-watching fMRI data, two infants completed two scans and two infants completed four scans each. The four infants who completed multiple movie-watching scans did so across multiple sessions (mean = 3.5 months between consecutive sessions, range = 1.4 − 6.3 months). Infant data were collected at the Brain Imaging Center in the Faculty of Arts and Sciences at Yale University (*N* = 26 movie-watching scans, *N* = 16 sleep scans) and at the Scully Center at Princeton University (*N* = 4 sleep scans). The study was approved by the Human Subjects Committee (HSC) at Yale University, and parents provided informed consent on behalf of their infants. Data used in the present paper were initially analyzed in [51] and openly accessible via DataDryad: https://doi.org/10.5061/dryad.nvx0k6dzf.

#### MRI data acquisition

Infant fMRI data and adult comparison data were collected using a previously validated procedure [62], described in brief here. All adult data and most infant data were acquired at the Brain Imaging Center in the Faculty of Arts and Sciences at Yale University using a Siemens Prisma 3T MRI using the bottom half of a 20-channel head coil. Functional images were acquired using a whole-brain T2* gradient-echo EPI sequence (*TR* = 2 s, *TE* = 30 ms, flip angle = 71, matrix = 64 × 64, slices = 34, resolution = 3mm isotropic, interleaved slice acquisition). For infants, anatomical images were collected as a T1 PETRA sequence (*TR*1 = 3.32ms, *TR*2 = 2250ms, *TE* = 0.07ms, flip angle = 6, matrix = 320 × 320, slices = 320 , resolution = 0.94 mm isotropic, radial slices = 30, 000), still with only the bottom half of the 20-channel head coil. For adults, anatomical images were collected including the top half of the 20-channel head coil as a T1 MPRAGE sequence (*TR* = 2300ms, *TE* = 2.96 ms, *TI* = 900ms, flip angle = 9, iPAT = 2, slices = 176, matrix = 256 × 256, resolution = 1.0mm isotropic). The remaining infant data were collected either on a Siemens Skyra 3T MRI at the Scully Center at Princeton University or a Siemens Prisma 3T MRI at the Magnetic Resonance Research Center (MRRC) at Yale University, with all the same acquisiton parameters except that the functional EPI sequence had slightly different parameters at these latter 2 sites (*TE* = 28 ms, slices = 36). All data collection procedures follow those outlined in prior publications [51, 62, 63].

#### MRI data processing

Both adult and infant data were preprocessed using a custom awake infant fMRI pipeline detailed in [62] and described here in brief. Data from infants were sometimes cleaved into pseudo-runs when another task was performed in the same functional run, and 3 burn-in volumes were discarded from the beginning of each run/pseudo-run. Volumes were realigned using slice-timing correction and timepoints with greater than 3 mm of translational motion were excluded and temporally interpolated so as to not bias linear detrending. Participants who had more than 96% of timepoints without motion were included, resulting in almost all timepoints being included during sleep runs (mean = 99.74% *±* 0.75; range across participants = 96.72 − 100%), infant movie runs (mean = 99.64% *±* 0.76; range across participants = 96.67 − 100%) , adult rest runs (all 100%), and adult movie runs (mean = 99.99% *±* 0.12%; range across participants = 98.89 − 100%). In analyses, we excluded motion confound timepoints. During infant movie runs, we also excluded timepoints when eyes were closed for the majority of movie frames. Data were linear detrended in time and spatially smoothed using a 5mm FWHM kernel before being z-scored within run/psuedo-run.

Anatomical images were aligned into standard space using the nonlinear alignment algorithm ANTs [64]. Infant images used an initial linear alignment using 12 DOF to align their anatomical data to an age-specific infant template in MNI space, followed by a nonlinear warping using diffeomorphic symmetric normalization [65]. Then, a predefined transformation (12 DOF) was used to linearly align between the infant template and the adult MNI152 template brain. Adult images used the same alignment procedure except that participants were directly aligned to the adult MNI152 template. Analyses were performed only in voxels which were included in the intersection of all brain masks for a given task and group.

#### Partly Cloudy dataset

##### MRI data acquisition

Full description about data acquisition can be found in the original publication [45]. In brief, whole-brain functional and anatomical images were collected on a Siemens Tim Trio 3T scanner using a 32-channel head coil. Children under 5 years used one of two custom 32-channel phased-array head coils made for small children. All other participants used a standard Siemens 32-channel head coil. Whole-brain fMRI data were acquired using a gradient-echo EPI sequence (TR = 2 s, TE = 30 ms, flip angle = 90, matrix = 64 × 64 , 32 interleaved slices).All functional data were upsampled in normalized space to 2 mm isotropic voxels. T1-weighted structural images were collected using a MPRAGE sequence (GRAPPA = 3, slices = 176, resolution 1 mm isotropic, FOV adult coil = 256 mm , FOV child coil = 192 mm).

##### MRI data processing

Data were preprocessed with fMRIPrep [61] and as described in [66]. In brief, fMRIPrep was used to perform coregistration, slice-time correction, motion correction, and nonlinear alignment to the MNI152 template brain. All functional data were subsequently upsampled in normalized space to 2 mm isotropic voxels. Timeseries data were detrended with regressors for (translation and rotation in the x, y, z dimensions), white matter, and cerebrospinal fluid, before smoothing with a 5 mm FWHM Gaussian kernel. Analyses were performed only in voxels which were included in the intersection of all participants’ brain masks.

#### Healthy Brain Network dataset

##### Participants

We used a subset of data from the Healthy Brain Network Biobank (HBN), a large study of children and adolescents aged 5–21 who undergo neuroimaging, cognitive, and clinical assessments across multiple sites in the New York City area. HBN aims to collect multimodal data from 10,000 individuals spanning broad range of clinical psychopathology. Data collection includes psychiatric, behavioral, cognitive, and lifestyle phenotypes and neuroimaging, biological, genetic, and behavioral recordings, to support psychiatric biomarker identification. In the full study, participants were excluded if they had serious neurological disorders preventing full participation in the study (e.g., nonverbal, chronic epilepsy); acute encephalopathy caused by injury or disease to the brain; known neurodegenerative disorders (e.g., ALS, Cerebral Palsy, Huntington’s disease, MS); hearing or visual impairment that is not corrected with devices; recent formal diagnosis (within past 6 months) of schizophrenia, schizoaffective disorder, or bipolar disorder; manic or psychotic episodes in the past 6 months without current, ongoing treatment; new onset (within past 3 months) of suicidality or homicidality without current, ongoing treatment; history of lifetime substance dependence requiring chemical replacement therapy; or acute intoxication at any point in the study visit. This study was approved by the Chesapeake Institutional Review board. Participants aged 18 and older provided written informed consent, and participants under 18 provided written assent while parents provided written consent. More information on these participants can be found at the original publication [52]. MRI data were collected at the following sites: Rutgers University Brain Imaging Center (RUBIC), Citigroup Biomedical Imaging Center (CBIC), City College of New York (CUNY), and a mobile scanner located in Staten Island (SI) (SI site was excluded in the present analyses).

In the current study, we focused on participants who completed both a movie-watching task (“The Present”) and at least one resting state run in the scanner. The Present is an emotionally-evocative short audiovisual animated movie (approx. 4 min) focusing on the relationship between a boy and a dog. The full protocol included two fixation rest scans each lasting 5 minutes (where participants were instructed to focus on a fixation cross), and we used only the first resting state scan here for analysis. At the time of analysis, the HBN Biobank contained some form of MRI data from 2,611 participants. We excluded participants who were scanned at the SI site, which used a mobile 1.5 T scanner as opposed to the 3T scanners at the other 3 sites. 683 of these participants had structural MRI datasets and complete scans for the movie-viewing task and at least one resting state scan. Participants were further excluded based upon quality control assessment, performed using fMRIprep (more details below), resulting in a total of 528 participants with usable data during both the movie-viewing and resting state scans. Of the total 528 participants (*N* = 204 female, *N* = 29 other, mean age = 12.32 *±* 3.5 years; range = 5.16 − 21.90 years), *N* = 282 were scanned at CBIC, *N* = 215 were scanned at RUBIC, and *N* = 31 were scanned at CUNY. Data were accessed via https://data.healthybrainnetwork.org/[52].

##### MRI data acquisition

MRI data were collected at one of three imaging centers. Images at the RUBIC site were acquired on a Siemens Tim Trio 3T scanner. Images at the CBIC and CUNY sites were collected on a Siemens Prisma 3T scanner. The three sites used identical fMRI parameters: 32-channel headcoils, whole-brain functional images collected in an interleaved fashion (60 slices per volume, 2.4mm isotropic resolution) using a T2* BOLD EPI sequence (*TR* = 800 ms, *TE* = 30 ms, flip angle = 31, FOV= 202 × 202 × 144) with a multiband acceleration factor of 6. T1w structural images were collected using a MPRAGE sequence (GRAPPA =, slices = 224, resolution = 0.8mm isotropic, TR = 2500ms, *TE* = 3.15m, flip angle= 8, FOV = 179 × 256 × 256 mm).

##### MRI data processing

Neuroimaging data were preprocessed using fMRIprep, a preprocessing tool based in Nipype that consists of both anatomical and functional pipelines. T1w images were corrected for intensity nonuniformity using N4BiasFieldCorrection [67] as distributed with ANTs [64] and skullstripped. Brain tissue segmentation of cerebrospinal fluid (CSF), white matter (WM), and gray matter (GM) was performed on the brain-extracted T1w using fast [68]. Brain surfaces were reconstructed using recon-all [69–71]. Volume-based spatial normalization to MNI152 standard space was performed with nonlinear registration with antsRegistration (ANTs 2.3.3) using a brain-extracted version of the T1w image. For BOLD preprocessing, head motion parameters were estimate with respect to a skull-stripped reference BOLD volume, including transformation matrices and rotation and translation parameters in the x,y,z directions. BOLD data were then slice-time corrected, resampled to native space, and co-registered to the T1w reference using boundary based registration with 6 degrees of freedom [72]. Confound timeseries were calculated based upon this preprocessed BOLD data, including framewise displacement (FD), DVARS, and global signals from CSF, WM, and the whole-brain mask. Participants whose structural or anatomical data were incomplete during fMRIprep or encountered errors related to MRIQC (quality control checker within fMRIprep) were excluded from subsequent analyses (resulting in 528 final participants in our sample). BOLD timeseries data were resampled into MNI152 standard space and later cleaned of nuisance signals by regressing the 6 motion parameters, WM, and CSF signals from the data and performing high-pass filtering, before smoothing with a 5 mm FWHM Gaussian kernel. Timeseries data were finally z-scored within voxel.

#### Benchmark dimensionality estimation methods

To compare the accuracy of our T-PHATE ID estimator, we employed a series of linear (e.g., principal components analysis) and nonlinear (e.g., maximum likelihood estimation) estimations on simulated manifolds and simulated fMRI data. These estimators were implemented in the `scikit-dimension` package [25]. All notation assumes the same input data and notation as for the T-PHATE algorithm: **X** ∈ ℝ^*T* ×*V*^ , where **X** denotes the data matrix for a given area, *T* is the number of time points, and *V* is the number of voxels within the area. ∈

##### Principal Component Analysis

As a linear baseline, we applied Principal Component Analysis (PCA) [73, 74]. After mean-centering the data, we computed the covariance matrix

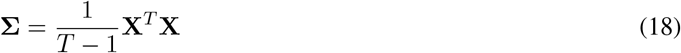

Let Λ_*T*_ denote the eigenvalues of **Σ**, where *λ*_1_ ≥ *λ*_2_ ≥ · · · ≥ *λ*_*T*_ . We computed the intrinsic dimensionality of **X** as the minimum number of eigenvalues required to account for a cumulative variance explained ratio of *θ*, where again *θ* = 0.9.

##### Local PCA (lPCA)

Local PCA estimates the intrinsic dimensionality by performing PCA locally within neighborhoods of each data point, then averaging the results. Rather than counting components above a variance threshold, lPCA uses a participation ratio (PR), which is a measure that quantifies the “effective number” of dimensions based on how uniformly variance is distributed across principal components. If the variance is concentrated in a few components (low PR), local dimensionality is low, whereas if the variance is spread across components (high PR), local dimensionality is high. The final ID estimate is the average PR across all local neighborhoods in the dataset.

Formally, for each data point **x**_**i**_, identify its k-nearest neighbors to form a local dataset 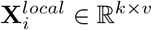 and compute the local covariance matrix:

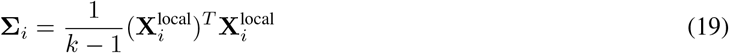

Eigendecompose **Σ**_*i*_ to obtain the eigenvalues {*λ*_*i*1_ ≥ *λ*_*i*2_ ≥ · · · ≥*λ*_*iT*_}. Then, compute the PR for the local neighborhood:

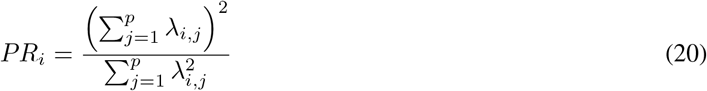

Finally, compute the intrinsic dimensionality estimate as the average PR across all points:

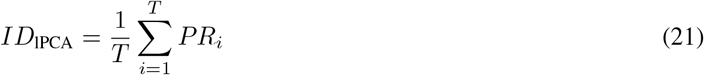

##### Maximum likelihood estimation (MLE)

The MLE method estimates intrinsic dimensionality based on the distribution of distances to nearest neighbors, assuming data are locally linear and uniformly distributed on a manifold [57]. The key insight is that in a *d*-dimensional space, the ratio of distances to the *k*-th and *j*-th nearest neighbors follows a predictable distribution that depends on *d*. By computing these distance ratios for each point’s neighborhood and finding the dimension that maximizes the likelihood of observing the data, MLE provides a local estimate of intrinsic dimensionality that can be averaged across all points.

Formally, for each data point **x**_*i*_, identify and compute and sort its distances to its *k*-nearest neighbors:

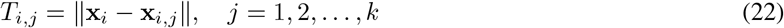

Then estimate the local MLE for each point *i* as:

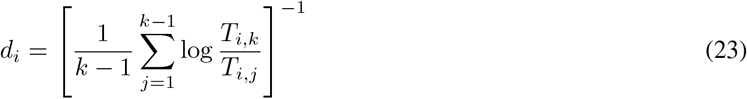

and the final ID estimation is:

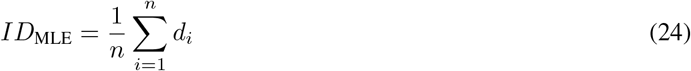

##### Minimum neighbor distance–Maximum likelihood (MiND_ML)

MiND_ML is a maximum likelihood estimator that adaptively accounts for manifold curvature and non-uniform sampling density. While standard MLE (as above) assumes the data are locally uniform on a flat manifold, MiND_ML builds upon MLE by incorporating correction terms that adjust the likelihood estimation based on local density variations and curvature, making it more robust in realistic scenarios. The method estimates intrinsic dimensionality locally around each point using the distribution of distances to its *k*-nearest neighbors, and aggregates local estimates to produce a global ID measure. [59].

Building upon the local MLE estimate for each data point *i* (*d*_*i*_) as in Equation 23, MiND_ML incorporates adaptive corrections for local geometry, where *C*_*i*_ represents local curvature and density corrections computed from the distribution of neighbor distances and *α*_*i*_ is an adaptive weighting parameter:

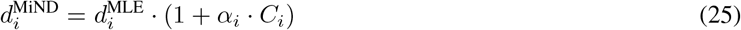

Finally, the final intrinsic dimensionality estimate is:

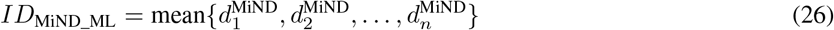

##### Dimensionality from angle and norm concentration (DANCo)

DANCo estimates intrinsic dimensionality by exploiting geometric properties of high-dimensional spaces: in truly high-dimensional data, random vectors become nearly orthogonal (angles concentrate near 90) and their norms become increasingly similar, but when data lies on a lower-dimensional manifold, these concentration effects are weaker. DANCo computes pairwise distances and angles between each data point and its neighbors, then analyzes the mean and variance of these quantities. By comparing observed concentration statistics to theoretical predictions for different dimensionalities, DANCo jointly leverages both angle and norm information to estimate the effective intrinsic dimension, making it robust to noise and non-uniform sampling [58].

Formally, for each data point **x**_*i*_, identify its *k*-nearest neighbors: 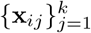. Then, compute vectors from **x**_*i*_ to its neighbors: **v**_*i,j*_ = **x**_*i,j*_ − **x**_*i*_. Calculate the distances and normalized direction vectors:

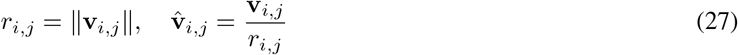

And the pairwise angles between neighbor vectors via their cosines:

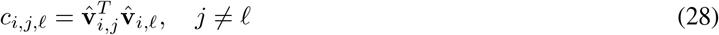

Then, estimate the local statistics (i.e., mean and variance) of the distances and angles:

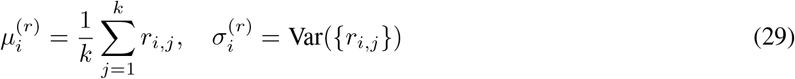

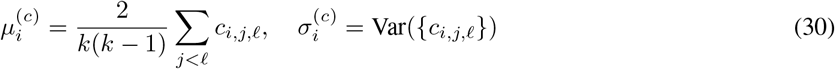

DANCo estimates the dimensionality for each point *i* by matching the observed statistics to theoretical models, where *f* represents an estimation procedure based on concentration theory:

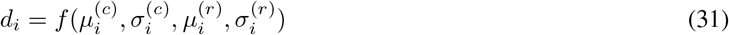

And the final estimation becomes:

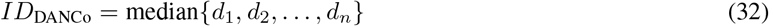

### Supplemental Tables

**Table S1:** Benchmark set of synthetic manifolds with know embedding and intrinsic dimensionality. Accessed from [25] and described by [53].

| Dataset Name | Intrinsic Dimensions<br>($ID_{true}$) | Embedding Dimensions<br>($ED$) | Description |
| --- | --- | --- | --- |
| **Helix** | 1 | 3 | 1D helix |
| **Nonlinear** | 6 | 36 | Nonlinear manifold |
| **Roll** | 2 | 3 | Swiss Roll |
| **Spiral** | 1 | 13 | 1D helix curve |
| **Paraboloid** | 3 | 12 | 3D paraboloid, nonlinearly embedded in a $3(3 + 1)D$ space according to a multivariate Burr distribution ($\alpha = 1$) |

**Table S2:** Participants included in each dataset and information about the tasks they performed. The 40 participants in the Narratives dataset all overlapped across tasks, as did the Healthy Brain Network and Rest/Movie – Adult. Under stimulus, A = auditory only, V = visual only, AV = audiovisual.

| Dataset | Stimulus | Task | $N$ | Age ($\mu \pm \sigma$) | %<br>Female | # TRs |
| --- | --- | --- | --- | --- | --- | --- |
| Narratives<br>[28] | Story (A) | Black | 40 | $23.5 \pm 7.9$ y | 72 | 550 |
| | | Bronx | 40 | $23.5 \pm 7.9$ y | 72 | 390 |
| | | Forgot | 40 | $23.5 \pm 7.9$ y | 72 | 574 |
| | | PiemanPNI | 40 | $23.5 \pm 7.9$ y | 72 | 294 |
| Adult Rest/Movie<br>[51] | Fixation | Rest | 12 | $21.4 \pm 3.4$ y | 58 | 152 |
| | Cartoon (V) | Aeronaut | 12 | $21.4 \pm 3.4$ y | 58 | 93 |
| Healthy Brain Network<br>[52] | Fixation | Rest | 528 | $12.3 \pm 3.5$ y | 44 | 375 |
| | Cartoon (AV) | The Present | 528 | $12.3 \pm 3.5$ y | 44 | 300 |
| Partly Cloudy<br>[45] | Cartoon (V) | Partly Cloudy | 155 | $10.5 \pm 8.1$ y | 54 | 168 |
| Infant Rest/Movie<br>[51] | None | Sleep | 20 | $11.2 \pm 5.1$ m | 73 | 61 |
| | Cartoon (V) | Aeronaut | 26 | $10.9 \pm 4.8$ m | 60 | 93 |

### Supplemental Figures

**Figure S1:**
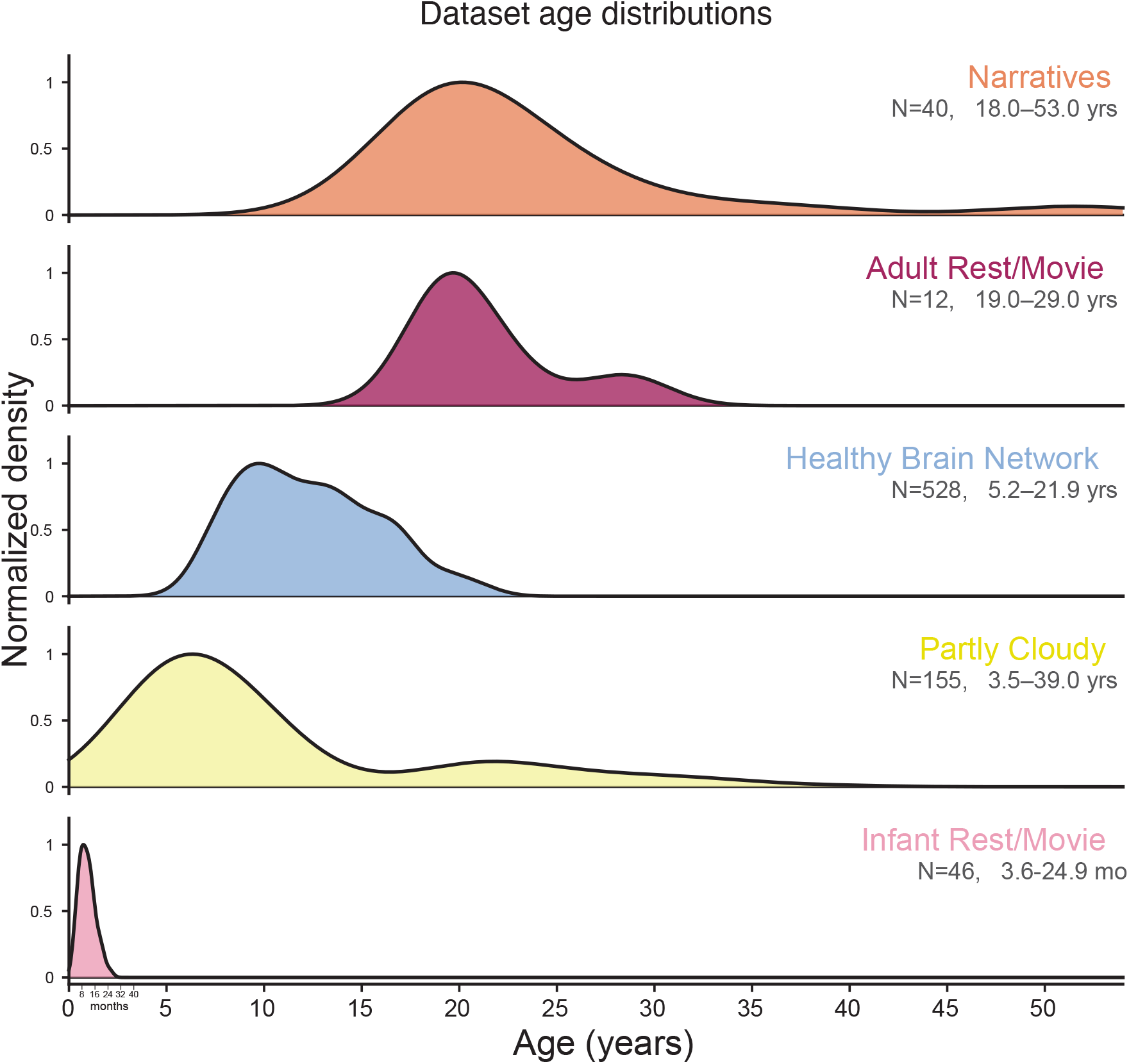
Distribution of participant ages for all datasets. Normalized age distributions are shown separately for each dataset. (A) The Narratives dataset includes adults aged 18-53 (*N* = 40). (B) The adult Rest/Movie dataset includes adults aged 19-29 years (*N* = 12). (C) The Healthy Brain Network (HBN) dataset includes children and adolescents aged 5-22 years (*N* = 528). (D) The Partly Cloudy dataset includes children ages 3-12 years (*N* = 122) and adults ages 18-39 years (*N* = 33). (E) The Infant Rest/Movie dataset includes infants age 3 - 25 months (*N* = 46), shown on a separate month scale (inset, left); sleeping and movie-watching scans are from separate sets of infant scanning sessions (see Table SS2).

**Figure S2:**
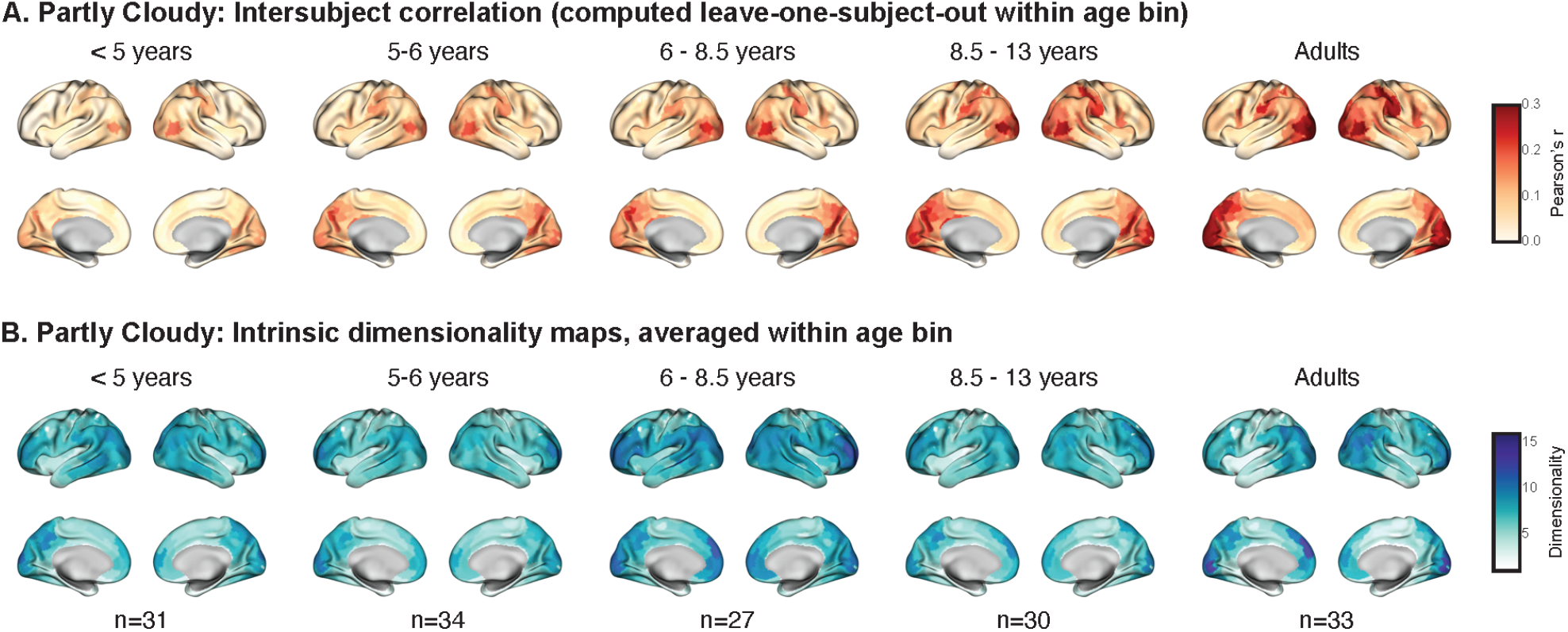
ISC and ID during movie-watching in the Partly Cloudy dataset. Participants were binned into < 5 years (*n* = 31), 5 − 6 years (*n* = 34), 6 − 8.5 years (*n* = 26), 8.5 − 13 years (*n* = 31), and adults (*n* = 33). (A) Group-averaged ISC maps for participants watching the silent cartoon movie “Partly Cloudy.” (B) Group-averaged ID maps for the same participants.

**Figure S3:**
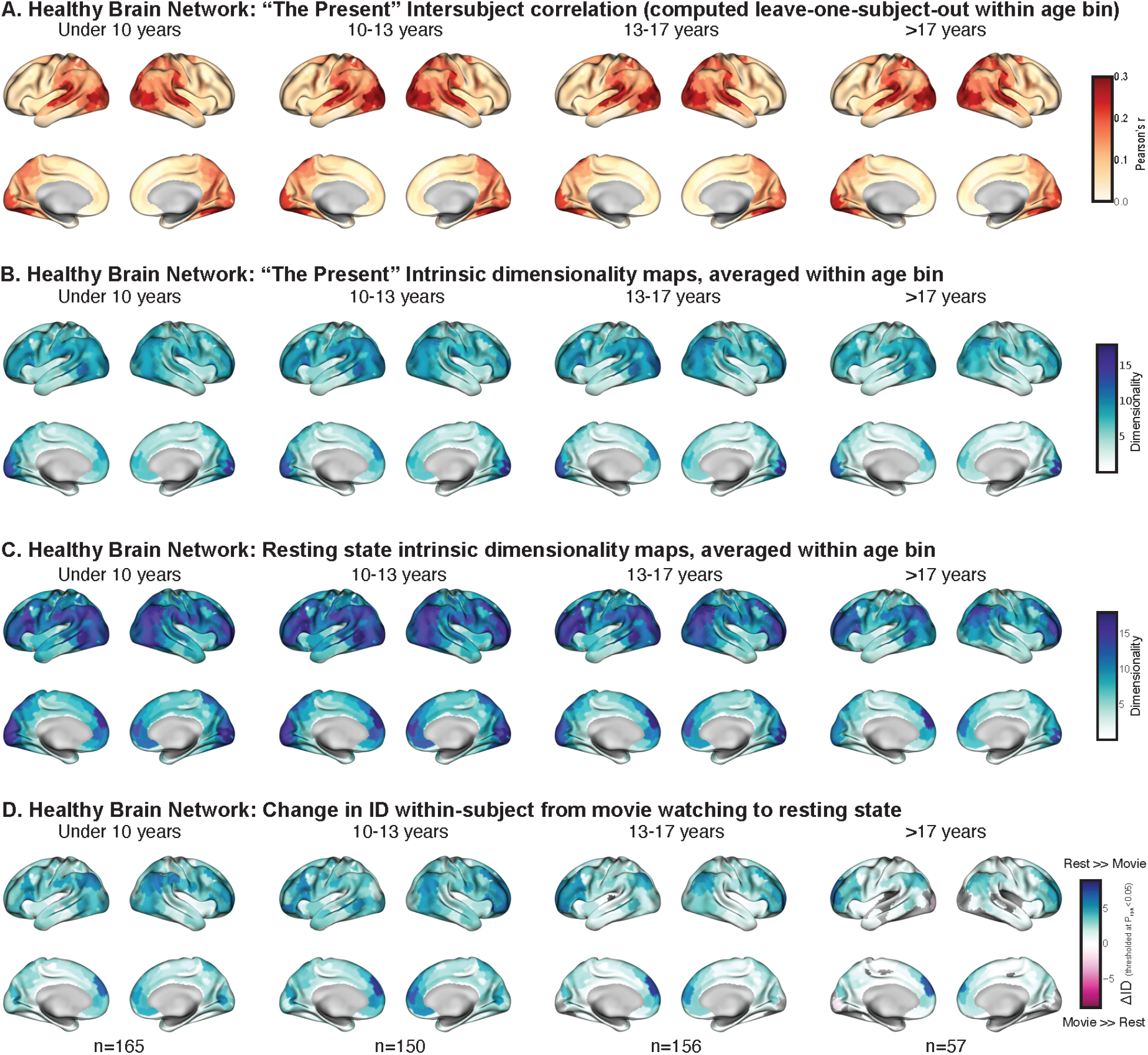
Group-averaged brain maps in the Healthy Brain Network (HBN) dataset. Participants were binned into < 10 years (*n* = 165), 10 − 13 years (*n* = 150), 13 − 17 years (*n* = 156), and ≥ 17 years (*n* = 57). (A) Group-averaged ISC maps for the HBN participants (ages 5-22 years, *N* = 528) watching the audiovisual cartoon movie “The Present.” Highest ISC was observed in visual and auditory cortices. Color reflects ISC magnitude. (B) Group-averaged ID maps as participants watched “The Present.” (C) Group-averaged ID maps for the same participants completing a fixation-rest run. (D) Group-averaged Δ*ID* maps for the same participants. Δ*ID* maps were computed within participant by subtracting a participant’s movie-watching ID map from their resting-state ID map. Statistical significance of these differences was computed across subjects, within age bin, using bootstrap resampling. Cooler colors indicate rest *>>* movie ID; warmer colors indicate movie *>>* rest ID. p-values were FDR corrected across parcels and visualized at *p*_*FDR*_ < 0.05.

**Figure S4:**
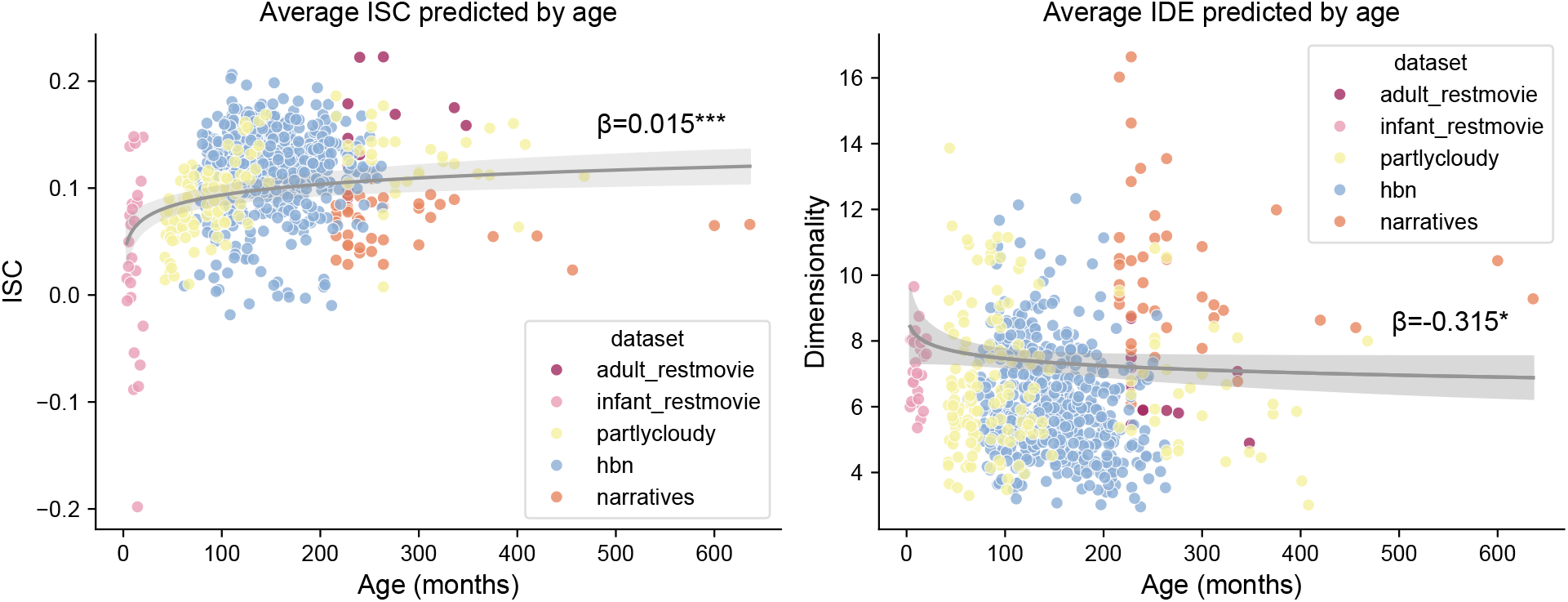
Mean ISC and ID as a function of age across datasets. Mean whole-brain ISC (left) or ID (right) as a function of age, pooled across the five datasets. Each point represents one participant; datasets are distinguished by color; line shows mixed-effects model fit, band shows the 95% CI of the model fit estimated with bootstrap resampling (10,000 iterations). (left) A mixed-effects model revealed a significant positive effect of log-transformed age on average whole-brain ISC (*β*_*age*_ = 0.015, 95% *CI* = [0.008, 0.021], *z* = 4.17, *p* = 0.00003), accounting for fixed effects of mean framewise displacement (FD) and sex and random effects of dataset. (Right) A mixed-effects model revealed a slight but significant negative effect of log-transformed age on average whole-brain ID (*β*_*age*_ = −0.315, 95% *CI* = [−0.626, −0.004], *z* = −1.986, *p* = −0.047), accounting for fixed effects of FD and sex and random effects of dataset. Together, these results indicate that while stimulus-driven responses strengthen with age, mean whole-brain ID decreases slightly, pointing to regionally specific rather than globally monotonic effects of development on dimensionality.

## References

1. Diedrichsen, J., Wiestler, T. & Ejaz, N. A multivariate method to determine the dimensionality of neural representation from population activity. NeuroImage 76, 225–235. ISSN: 1053-8119. https://www.sciencedirect.com/science/article/pii/S1053811913002176 (2026) (Aug. 2013).

2. Ahlheim, C. & Love, B. C. Estimating the functional dimensionality of neural representations. NeuroImage 179, 51–62. ISSN: 1053-8119. https://www.sciencedirect.com/science/article/pii/S1053811918305226 (2025) (Oct. 2018).

3. Weaver, N. J., Faskowitz, J., Betzel, R. F. & Lynn, C. W. Quantifying the compressibility of the human brain. Proceedings of the National Academy of Sciences 123, e2531115123. https://www.pnas.org/doi/10.1073/pnas.2531115123 (2026) (Jan. 2026).

4. Camastra, F. & Staiano, A. Intrinsic dimension estimation: Advances and open problems. Information Sciences 328, 26–41. ISSN: 0020-0255. https://www.sciencedirect.com/science/article/pii/S0020025515006179 (2026) (Jan. 2016).

5. Jazayeri, M. & Ostojic, S. Interpreting neural computations by examining intrinsic and embedding dimensionality of neural activity. Current Opinion in Neurobiology.Computational Neuroscience 70, 113–120. ISSN: 0959-4388. https://www.sciencedirect.com/science/article/pii/S0959438821000933 (2024) (Oct. 2021).

6. Gao, P. & Ganguli, S. On simplicity and complexity in the brave new world of large-scale neuroscience. Current Opinion in Neurobiology. Large-Scale Recording Technology (32) 32, 148–155. ISSN: 0959-4388. https://www.sciencedirect.com/science/article/pii/S0959438815000768 (2026) (June 2015).

7. Fusi, S., Miller, E. K. & Rigotti, M. Why neurons mix: high dimensionality for higher cognition. Current Opinion in Neurobiology. Neurobiology of cognitive behavior 37, 66–74. ISSN: 0959-4388. https://www.sciencedirect.com/science/article/pii/S0959438816000118 (2025) (Apr. 2016).

8. Rigotti, M. et al. The importance of mixed selectivity in complex cognitive tasks. en. Nature 497. Number: 7451, 585–590. ISSN: 1476-4687. https://www.nature.com/articles/nature12160 (2023) (May 2013).

9. Cunningham, J. P. & Yu, B. M. Dimensionality reduction for large-scale neural recordings. en. Nature Neuroscience 17. Number: 11, 1500–1509. ISSN: 1546-1726. https://www.nature.com/articles/nn.3776 (2020) (Nov. 2014).

10. Tang, E. et al. Effective learning is accompanied by high-dimensional and efficient representations of neural activity. en. Nature Neuroscience 22. Number: 6, 1000–1009. ISSN: 1546-1726. https://www.nature.com/articles/s41593-019-0400-9 (2020) (June 2019).

11. Kikumoto, A., Bhandari, A., Shibata, K. & Badre, D. A transient high-dimensional geometry affords stable conjunctive subspaces for efficient action selection. en. Nature Communications 15, 8513. ISSN: 2041-1723. https://www.nature.com/articles/s41467-024-52777-6 (2026) (Oct. 2024).

12. Sheng, J. et al. Higher-dimensional neural representations predict better episodic memory. Science Advances 8, eabm3829. https://www.science.org/doi/10.1126/sciadv.abm3829 (2022) (Apr. 2022).

13. Areshenkoff, C. et al. Neural excursions from manifold structure explain patterns of learning during human sensorimotor adaptation. eng. eLife 11, e74591. ISSN: 2050-084X (Apr. 2022).

14. Gallego, J. A. et al. Cortical population activity within a preserved neural manifold underlies multiple motor behaviors. eng. Nature Communications 9, 4233. ISSN: 2041-1723 (Oct. 2018).

15. Perich, M. G., Narain, D. & Gallego, J. A. A neural manifold view of the brain. en. Nature Neuroscience 28, 1582–1597. ISSN: 1546-1726. https://www.nature.com/articles/s41593-025-02031-z (2025) (Aug. 2025).

16. Ganguli, S. Measuring the dimensionality of behavior. Proceedings of the National Academy of Sciences 119, e2205791119. https://www.pnas.org/doi/abs/10.1073/pnas.2205791119 (2025) (Oct. 2022).

17. Churchland, M. M. et al. Stimulus onset quenches neural variability: a widespread cortical phenomenon. en. Nature Neuroscience 13. Number: 3, 369–378. ISSN: 1546-1726. https://www.nature.com/articles/nn.2501 (2023) (Mar. 2010).

18. Han, C. & Bonner, M. F. High-dimensional structure underlying individual differences in naturalistic visual experience arXiv:2505.12653 [q-bio]. Dec. 2025. http://arxiv.org/abs/2505.12653 (2026).

19. Chen, Z. & Bonner, M. F. Universal dimensions of visual representation. Science Advances 11, eadw7697. https://www.science.org/doi/full/10.1126/sciadv.adw7697 (2026) (July 2025).

20. Badre, D., Bhandari, A., Keglovits, H. & Kikumoto, A. The dimensionality of neural representations for control. Current Opinion in Behavioral Sciences.Computational cognitive neuroscience 38, 20–28. ISSN: 2352-1546. https://www.sciencedirect.com/science/article/pii/S2352154620301042 (2026) (Apr. 2021).

21. Busch, E. L. et al. Multi-view manifold learning of human brain-state trajectories. en. Nature Computational Science, 1–14. ISSN: 2662-8457. https://www.nature.com/articles/s43588-023-00419-0 (2023) (Mar. 2023).

22. Olszowy, W., Aston, J., Rua, C. & Williams, G. B. Accurate autocorrelation modeling substantially improves fMRI reliability. Nature communications 10, 1220 (2019).

23. Stroud, J. P. et al. Effects of noise and metabolic cost on cortical task representations. eLife 13, RP94961. ISSN: 2050-084X. https://pmc.ncbi.nlm.nih.gov/articles/PMC11750133/ (2026).

24. De, A. & Chaudhuri, R. Common population codes produce extremely nonlinear neural manifolds. Proceedings of the National Academy of Sciences 120, e2305853120. https://www.pnas.org/doi/full/10.1073/pnas.2305853120 (2025) (Sept. 2023).

25. Bac, J., Mirkes, E. M., Gorban, A. N., Tyukin, I. & Zinovyev, A. Scikit-Dimension: A Python Package for Intrinsic Dimension Estimation. en. Entropy 23. Number: 10, 1368. ISSN: 1099-4300. https://www.mdpi.com/1099-4300/23/10/1368 (2023) (Oct. 2021).

26. Schaefer, A. et al. Local-Global Parcellation of the Human Cerebral Cortex from Intrinsic Functional Connectivity MRI. Cerebral Cortex (New York, NY) 28, 3095–3114. ISSN: 1047-3211. https://www.ncbi.nlm.nih.gov/pmc/articles/PMC6095216/(2024) (Sept. 2018).

27. Moon, K. R. et al. Visualizing structure and transitions in high-dimensional biological data. en. Nature Biotechnology 37. Number: 12, 1482–1492. ISSN: 1546-1696. https://www.nature.com/articles/s41587-019-0336-3 (2020) (Dec. 2019).

28. Nastase, S. A. et al. The “Narratives” fMRI dataset for evaluating models of naturalistic language comprehension. en. Scientific Data 8. Number: 1, 250. ISSN: 2052-4463. https://www.nature.com/articles/s41597-021-01033-3 (2023) (Sept. 2021).

29. Hasson, U., Nir, Y., Levy, I., Fuhrmann, G. & Malach, R. Intersubject Synchronization of Cortical Activity During Natural Vision. Science 303, 1634–1640. https://www.science.org/doi/10.1126/science.1089506 (2026) (Mar. 2004).

30. Nastase, S. A., Gazzola, V., Hasson, U. & Keysers, C. Measuring shared responses across subjects using intersubject correlation. Social Cognitive and Affective Neuroscience 14, 667–685. ISSN: 1749-5016. 10.1093/scan/nsz037 (2026) (Aug. 2019).

31. Johnson, M. H. Interactive Specialization: A domain-general framework for human functional brain development? Developmental Cognitive Neuroscience 1, 7–21. ISSN: 1878-9293. https://www.sciencedirect.com/science/article/pii/S1878929310000046 (2022) (Jan. 2011).

32. Morfoisse, T. et al. Initial organization and progressive expansion of the math-responsive brain network during the first school years. Proceedings of the National Academy of Sciences 123, e2602515123 (2026).

33. Bethlehem, R. A. et al. Brain charts for the human lifespan. Nature 604, 525–533 (2022).

34. Mazzucato, L., Fontanini, A. & La Camera, G. Stimuli Reduce the Dimensionality of Cortical Activity. Frontiers in Systems Neuroscience 10. ISSN: 1662-5137. https://www.frontiersin.org/articles/10.3389/fnsys.2016.00011 (2023) (2016).

35. Owen, L. L. W. & Manning, J. R. High-level cognition is supported by information-rich but compressible brain activity patterns. Proceedings of the National Academy of Sciences 121, e2400082121. https://www.pnas.org/doi/abs/10.1073/pnas.2400082121 (2026) (Aug. 2024).

36. Stringer, C., Pachitariu, M., Steinmetz, N., Carandini, M. & Harris, K. D. High-dimensional geometry of population responses in visual cortex. en. Nature 571. Number: 7765, 361–365. ISSN: 1476-4687. https://www.nature.com/articles/s41586-019-1346-5 (2023) (July 2019).

37. Stringer, C. et al. Spontaneous behaviors drive multidimensional, brainwide activity. Science 364, eaav7893. https://www.science.org/doi/10.1126/science.aav7893 (2023) (Apr. 2019).

38. Nastase, S. A. et al. Attention Selectively Reshapes the Geometry of Distributed Semantic Representation. Cerebral Cortex 27, 4277–4291. ISSN: 1047-3211. 10.1093/cercor/bhx138 (2023) (Aug. 2017).

39. Song, H., Park, J. & Rosenberg, M. D. Understanding cognitive processes across spatial scales of the brain. English. Trends in Cognitive Sciences 0. ISSN: 1364-6613, 1879-307X. https://www.cell.com/trends/cognitive-sciences/abstract/S1364-6613(24)00252-3 (2024) (Nov. 2024).

40. Ostojic, S. & Fusi, S. Computational role of structure in neural activity and connectivity. English. Trends in Cognitive Sciences 28, 677–690. ISSN: 1364-6613, 1879-307X. https://www.cell.com/trends/cognitive-sciences/abstract/S1364-6613(24)00056-1 (2025) (July 2024).

41. Wei, X.-X. & Woodford, M. Representational geometry explains puzzling error distributions in behavioral tasks. Proceedings of the National Academy of Sciences 122, e2407540122. https://www.pnas.org/doi/10.1073/pnas.2407540122 (2026) (Jan. 2025).

42. Pospisil, D. A. & Pillow, J. W. Revisiting the high-dimensional geometry of population responses in the visual cortex. Proceedings of the National Academy of Sciences 122, e2506535122. https://www.pnas.org/doi/10.1073/pnas.2506535122 (2026) (Nov. 2025).

43. Jacoby, N., Bruneau, E., Koster-Hale, J. & Saxe, R. Localizing Pain Matrix and Theory of Mind networks with both verbal and non-verbal stimuli. NeuroImage 126, 39–48. ISSN: 1053-8119. https://www.sciencedirect.com/science/article/pii/S1053811915010472 (2026) (Feb. 2016).

44. Samara, A., Eilbott, J., Margulies, D. S., Xu, T. & Vanderwal, T. Cortical gradients during naturalistic processing are hierarchical and modality-specific. en. NeuroImage 271, 120023. ISSN: 1053-8119. https://www.sciencedirect.com/science/article/pii/S1053811923001696 (2023) (May 2023).

45. Richardson, H., Lisandrelli, G., Riobueno-Naylor, A. & Saxe, R. Development of the social brain from age three to twelve years. en. Nature Communications 9, 1027. ISSN: 2041-1723. https://www.nature.com/articles/s41467-018-03399-2 (2025) (Mar. 2018).

46. Park, B.-Y. et al. An expanding manifold in transmodal regions characterizes adolescent reconfiguration of structural connectome organization. eng. eLife 10, e64694. ISSN: 2050-084X (Mar. 2021).

47. Wójcik, M. J. et al. Learning shapes neural geometry in the prefrontal cortex en. ISSN: 2692-8205 Pages: 2023.04.24.538054 Section: New Results. Jan. 2026. https://www.biorxiv.org/content/10.1101/2023.04.24.538054v3 (2026).

48. Bartolo, R., Saunders, R. C., Mitz, A. R. & Averbeck, B. B. Dimensionality, information and learning in prefrontal cortex. en. PLOS Computational Biology 16, e1007514. ISSN: 1553-7358. https://journals.plos.org/ploscompbiol/article?id=10.1371/journal.pcbi.1007514 (2026) (Apr. 2020).

49. Rigotti, M. & Fusi, S. Estimating the dimensionality of neural responses with fMRI Repetition Suppression arXiv:1605.03952 [q-bio]. May 2016. http://arxiv.org/abs/1605.03952 (2025).

50. Luczak, A., Barthó, P. & Harris, K. D. Spontaneous Events Outline the Realm of Possible Sensory Responses in Neocortical Populations. English. Neuron 62, 413–425. ISSN: 0896-6273. https://www.cell.com/neuron/abstract/S0896-6273(09)00237-2 (2026) (May 2009).

51. Yates, T. S., Ellis, C. T. & Turk-Browne, N. B. Functional networks in the infant brain during sleep and wake states. Cerebral Cortex 33, 10820–10835. ISSN: 1047-3211. https://doi.org/10.1093/cercor/bhad327 (2025) (Nov. 2023).

52. Alexander, L. M. et al. An open resource for transdiagnostic research in pediatric mental health and learning disorders. en. Scientific Data 4, 170181. ISSN: 2052-4463. https://www.nature.com/articles/sdata2017181 (2026) (Dec. 2017).

53. Campadelli, P., Casiraghi, E., Ceruti, C. & Rozza, A. Intrinsic Dimension Estimation: Relevant Techniques and a Benchmark Framework. en. Mathematical Problems in Engineering 2015. _eprint: https://onlinelibrary.wiley.com/doi/pdf/10.1155/2015/759567, 759567. ISSN: 1563-5147. https://onlinelibrary.wiley.com/doi/abs/10.1155/2015/759567 (2025) (2015).

54. Busch, E. L., Turk-Browne, N. B. & Baskin-Sommers, A. Revamping neuroimaging analysis to reveal biomarkers of adolescent mental health. Nature Mental Health, 1–13 (2026).

55. Vos de Wael, R. et al. BrainSpace: a toolbox for the analysis of macroscale gradients in neuroimaging and connectomics datasets. en. Communications Biology 3, 103. ISSN: 2399-3642. https://www.nature.com/articles/s42003-020-0794-7 (2026) (Mar. 2020).

56. Markello, R. D. et al. neuromaps: structural and functional interpretation of brain maps. en. Nature Methods 19, 1472–1479. ISSN: 1548-7105. https://www.nature.com/articles/s41592-022-01625-w (2026) (Nov. 2022).

57. Levina, E. & Bickel, P. Maximum Likelihood Estimation of Intrinsic Dimension in Advances in Neural Information Processing Systems 17 (MIT Press, 2004). https://papers.nips.cc/paper_files/paper/2004/hash/74934548253bcab8490ebd74afed7031-Abstract.html (2026).

58. Ceruti, C. et al. DANCo: An intrinsic dimensionality estimator exploiting angle and norm concentration. Pattern Recognition 47, 2569–2581. ISSN: 0031-3203. https://www.sciencedirect.com/science/article/pii/S003132031400065X (2026) (Aug. 2014).

59. Rozza, A., Lombardi, G., Ceruti, C., Casiraghi, E. & Campadelli, P. Novel high intrinsic dimensionality estimators. en. Machine Learning 89, 37–65. ISSN: 1573-0565. https://doi.org/10.1007/s10994-012-5294-7 (2025) (Oct. 2012).

60. Busch, E., Bhaskar, D. & Horvát, S. KrishnaswamyLab/TPHATE: v1.2.1 version v1.2.1. Nov. 2025. 10.5281/zenodo.17583910.

61. Esteban, O. et al. fMRIPrep: a robust preprocessing pipeline for functional MRI. eng. Nature Methods 16, 111–116.ISSN: 1548-7105 (Jan. 2019).

62. Ellis, C. T. et al. Re-imagining fMRI for awake behaving infants. en. Nature Communications 11, 4523. ISSN: 2041-1723. https://www.nature.com/articles/s41467-020-18286-y (2026) (Sept. 2020).

63. Yates, T. S. et al. Neural event segmentation of continuous experience in human infants. Proceedings of the National Academy of Sciences 119, e2200257119. https://www.pnas.org/doi/10.1073/pnas.2200257119 (2026) (Oct. 2022).

64. Avants, B. B. et al. A reproducible evaluation of ANTs similarity metric performance in brain image registration. eng. NeuroImage 54, 2033–2044. ISSN: 1095-9572 (Feb. 2011).

65. Fonov, V., Evans, A., McKinstry, R., Almli, C. & Collins, D. Unbiased nonlinear average age-appropriate brain templates from birth to adulthood. NeuroImage. Organization for Human Brain Mapping 2009 Annual Meeting 47, S102. ISSN: 1053-8119. https://www.sciencedirect.com/science/article/pii/S1053811909708845 (2026) (July 2009).

66. Yates, T. S., Ellis, C. T. & Turk-Browne, N. B. Emergence and organization of adult brain function throughout child development. NeuroImage 226, 117606. ISSN: 1053-8119. https://www.sciencedirect.com/science/article/pii/S1053811920310910 (2026) (Feb. 2021).

67. Tustison, N. J. et al. N4ITK: Improved N3 Bias Correction. IEEE Transactions on Medical Imaging 29, 1310–1320. ISSN: 1558-254X. https://ieeexplore.ieee.org/document/5445030 (2026) (June 2010).

68. Zhang, Y., Brady, M. & Smith, S. Segmentation of brain MR images through a hidden Markov random field model and the expectation-maximization algorithm. IEEE Transactions on Medical Imaging 20, 45–57. ISSN: 1558-254X. https://ieeexplore.ieee.org/document/906424 (2026) (Jan. 2001).

69. Fischl, B. FreeSurfer. NeuroImage. 20 YEARS OF fMRI 62, 774–781. ISSN: 1053-8119. https://www.sciencedirect.com/science/article/pii/S1053811912000389 (2022) (Aug. 2012).

70. Dale, A. M., Fischl, B. & Sereno, M. I. Cortical surface-based analysis. I. Segmentation and surface reconstruction. eng. NeuroImage 9, 179–194. ISSN: 1053-8119 (Feb. 1999).

71. Fischl, B., Sereno, M. I., Tootell, R. B. & Dale, A. M. High-resolution intersubject averaging and a coordinate system for the cortical surface. en. Human Brain Mapping 8. _eprint: https://onlinelibrary.wiley.com/doi/pdf/10.1002/%28SICI%291097-0193%281999%298%3A4%3C272%3A%3AAID-HBM10%3E3.0.CO%3B2-4, 272–284. ISSN: 1097-0193. https://onlinelibrary.wiley.com/doi/abs/10.1002/%28SICI%291097-0193%281999%298%3A4%3C272%3A%3AAID-HBM10%3E3.0.CO%3B2-4 (2022) (1999).

72. Greve, D. N. & Fischl, B. Accurate and robust brain image alignment using boundary-based registration. NeuroImage 48, 63–72. ISSN: 1053-8119. https://www.sciencedirect.com/science/article/pii/S1053811909006752 (2024) (Oct. 2009).

73. Pearson, K. LIII. On lines and planes of closest fit to systems of points in space. The London, Edinburgh, and Dublin Philosophical Magazine and Journal of Science 2. _eprint: 10.1080/14786440109462720, 559–572. ISSN: 1941-5982. https://doi.org/10.1080/14786440109462720 (2026) (Nov. 1901).

74. Hotelling, H. Analysis of a complex of statistical variables into principal components. Journal of Educational Psychology 24. Place: US, 417–441. ISSN: 1939-2176 (1933).

